# Calcium signaling is a universal carbon source signal transducer and effects an ionic memory of past carbon sources

**DOI:** 10.1101/2025.02.11.637612

**Authors:** Kobi Simpson-Lavy, Martin Kupiec

**Affiliations:** The Shmunis School of Biomedicine and Cancer Research, The George S. Wise Faculty of Life Sciences, Tel Aviv University, Ramat Aviv 69978, Israel

**Keywords:** Calcium, Yeast, Snf1, Carbon Metabolism, *ADH2*, *ZWF1*

## Abstract

Glucose is the preferred carbon source for most cells. However, cells may encounter other carbon sources that can be utilized. How cells match their metabolic gene expression to their carbon source, beyond a general glucose repressive system (catabolite repression), remains little understood. By studying the effect of up to seven different carbon sources on Snf1 phosphorylation and on the expression of downstream regulated genes, we searched for the mechanism that identifies carbon sources. We found that the glycolysis metabolites glucose-6-phosphate (G6P) and glucose-1-phosphate (G1P) play a central role in the adaptation of gene expression to different carbon sources. The ratio of G1P and G6P activates an analogue calcium signaling via the proton-exporter Pma1, to regulate downstream genes. The signaling pathway bifurcates with calcineurin reducing *ADH2* (alcohol dehydrogenase) expression and with Cmk1 increasing *ZWF1* (glucose-6-phosphate dehydrogenase) expression. Furthermore, calcium signaling is not only regulated by the present carbon source; it is also regulated by past carbon sources. We were able to manipulate this **ionic memory** mechanism to obtain high expression of *ZWF1* in media containing galactose. Our findings provide a **universal** mechanism by which cells respond to all carbon sources.

## Introduction

The availability of different carbon sources to an independent single celled organism affects that cell’s metabolism, growth, and resistance to adversity. Although the sensing and response to glucose or fructose (catabolite repression) has been extensively investigated (reviewed in [1]), how other carbon sources regulate cellular processes is poorly understood (except as a general no-glucose condition). Even within the “glucose response” body of research, a majority of research has been into the activation of PKA by glucose to inhibit stress-responsive genes and the inhibition of Snf1 by glucose to explain glucose-repression of respiratory genes. Other elements of glucose signaling remain less characterized; such as the regulation by Snf1 of the glucose-sensor activation of the Yck1/2 kinases [2,3] to regulate hexose-transporter expression and protein degradation [4–6] , the still mysterious activation of the plasma membrane ATPase Pma1 pump, which consumes 50% of a glucose-grown cell’s ATP [7], and calcium signaling [extensively reviewed in [8]]. Furthermore, aside from the induction of hexose transporters by glucose via external sensing by Rgt2 and Snf3 [4–6], most known glucose-regulated signaling is repressive, and a mechanism of how glucose <u>induces</u> gene expression remains elusive.

The classical catabolite repression model of carbon-source gene regulation holds that glucose/fructose/mannose suppress metabolism and metabolism-related gene expression pertaining to other carbon sources, and this is regulated by PKA activation in response to glucose [9] and Snf1 activation in the absence of glucose [10]. Snf1 regulation of catabolite repression has been much studied, with a disproportionate focus on the Mig1 repressor which is inhibited by Snf1 phosphorylation and disinhibited by Glc7-Reg1 dephosphorylation (e.g. [11,12]). However, much of the literature compares glucose to one other carbon source, usually galactose or glycerol, with little research comparing non-glucose carbon sources, which is somewhat surprising given that sucrose and maltose are the most important carbon sources commercially. Some carbon sources such as sucrose or glycerol have no known sensing mechanism, and although there are some transcription factors that respond to ethanol (e.g. Etp1 [13], Ert1 [14]) there is no known ethanol sensor. Only for the relationship between acetate and ethanol we understand the molecular mechanism: the acetate anion binds directly to the Haa1 transcription factor [15] and this activation of Haa1 represses *ADH2* expression to lower ethanol catabolism [16]. This also indicates the existence of more complex regulation within the set of non-glucose (fructose, mannose) carbon sources.

Glucose is transported into the cell by seventeen hexose transporters (Hxt1-17), whereupon it is phosphorylated by Hxk2 (and also by Hxk1 and Glk1) [17] . There are three fates of Glucose-6-phophate (G6P) (**Figure S1A**): **a)** Isomerization by Pgi1 to fructose-6-phosphate followed by a second phosphorylation by Phosphofructokinase (Pfk1-Pfk2) and glycolytic catabolism to pyruvate. **b)** Entry into the pentose-phosphate pathway by Zwf1 [18]; and **c)** conversion to Glucose-1-phosphate (G1P) by phosphoglucose mutatases (PGMs: mainly Pgm1 and Pgm2) [19]. G1P is converted to UDP-glucose by Ugp1, which is the precursor of the storage carbohydrates glycogen and trehalose, and of the β-glycans and glucosamines that comprise the cell wall. Galactose is metabolized to G1P by the Leloir pathway; and is converted to G6P by PGM to enter glycolysis (**Figure S1A**).

In *S. cerevisiae,* import of extracellular calcium is a response to depletion of internal calcium stores (primarily the vacuole) [20–22]. Calcium ions are imported into the cytoplasm from the external medium by the Mid1-Cch1 channel [23]. There are five known intracellular calcium pumps that sequester calcium into internal stores to maintain a cytoplasmic calcium concentration of 70-80nM in glucose-grown cells [24] (with a total calcium concentration of 1.5-4mM [24–26]). Calcium is released from these stores into the cytoplasm following exposure to glucose [27], cold stress [28], alkali stress [29], hypo-osmotic shock [22,30], toxic metals [28], [31],[32], and mating pheromone [33]. Exposure to glucose causes the release of calcium into the cytoplasm through the vacuolar Yvc1 channel [30] and other as yet unidentified channels [27]. This causes a rapid and **T**ransient **E**levation of **C**ytosolic **C**alcium levels (TECC) [34,35] that occurs 40-60 seconds after addition of glucose [19]. The same pumps then re-sequester the calcium in the stores [34,36], [37].

The TECC response is dependent upon glucose import and phosphorylation by hexokinases [19], and a strain deficient for all hexose and maltose transporters fails to produce a TECC pulse upon exposure to glucose [19]. Moreover, an *hxk1Δ hxk2Δ glk1Δ* yeast strain, despite being capable of importing glucose, cannot phosphorylate it [17] or produce a TECC response upon exposure to glucose [38]. Following addition of glucose, Glucose-6-phosphate (G6P) and Glucose-1-phosphate (G1P) levels increase after 30 seconds, preceding the TECC response (40-60 seconds after addition) [19]. The TECC response following glucose addition is delayed and diminished in *pgm2Δ* cells (which exhibit lower G1P levels) [19]. In summary, the balance between G1P and G6P regulates calcium accumulation: galactose-grown *pgm2Δ* cells (which exhibit high G1P) accumulate calcium; this can be suppressed by deletion of *PFK2* (thus increasing G6P) [39]. Inhibition of PGM activity by 10mM LiCl has the same effects on metabolite levels and calcium accumulation as *PGM2* deletion [40]. How the ratio between G6P and G1P is converted into a calcium signal remains to be determined. Once released into the cytoplasm, calcium acts as a second messenger to activate (via Calmodulin) several kinases and the phosphatase Calcineurin, whose activation of the Crz1 transcription factor has been extensively investigated [reviewed in [41]].

In this work we have examined up to seven different carbon sources for their effects on Snf1 phosphorylation and expression of downstream regulated genes. We find that calcium signaling provides a universal signaling mechanism in response to all carbon sources and regulates carbon source-dependent genes. In addition, calcium accumulation constitutes an ionic memory of prior carbon source exposure that modulates the response to fresh carbon source feeding.

## Results

### Snf1 phosphorylation levels do not correspond with *ADH2* expression

Snf1 activity is required to switch from hexose fermentation to the respiration of poor carbon sources such as ethanol. By activating Adr1, the expression of *ADH2* [encoding Alcohol Dehydrogenase 2; which catalyzes the first step of ethanol catabolism (**Figure 1A**)], commonly used as a reporter for Snf1 activity, is induced. In the presence of glucose (or fructose), phosphorylation of Snf1 at T210 (which is usually taken as being indicative of Snf1 activation) is low (**Figure 1B**). After shifting the cells to other carbon sources for 15 minutes there is a moderate amount of phosphorylation in the presence of sucrose (**Figure 1B, 1C**), despite Snf1 being required for growth with sucrose as the carbon source (**Figure S1A**). On other carbon sources (or in the absence of a carbon source) T210 phosphorylation is high, albeit less so in ethanol containing media (**Figure 1B, 1C**). We also examined Snf1 phosphorylation after 3 hours of growth in alternative carbon sources (in this experiment native genomic Snf1 was detected in wild-type cells) (**Figure S1B, C**). After 3 hours of growth in sucrose, Snf1 phosphorylation is similar to that of glucose-grown cells probably due to invertase activity, and the Snf1 phosphorylation of maltose grown cells is lower compared with the 15-minute time-point. Snf1 phosphorylation in galactose- and glycerol-grown cells is the same after three hours as ethanol- (or no carbon source-) grown cells (**Figure S1B, C**). In contrast, the expression of *ADH2* is markedly lower in galactose and glycerol compared with ethanol, and is absent entirely in sucrose containing media (**Figure 1D**) [16] .

**Figure 1.**
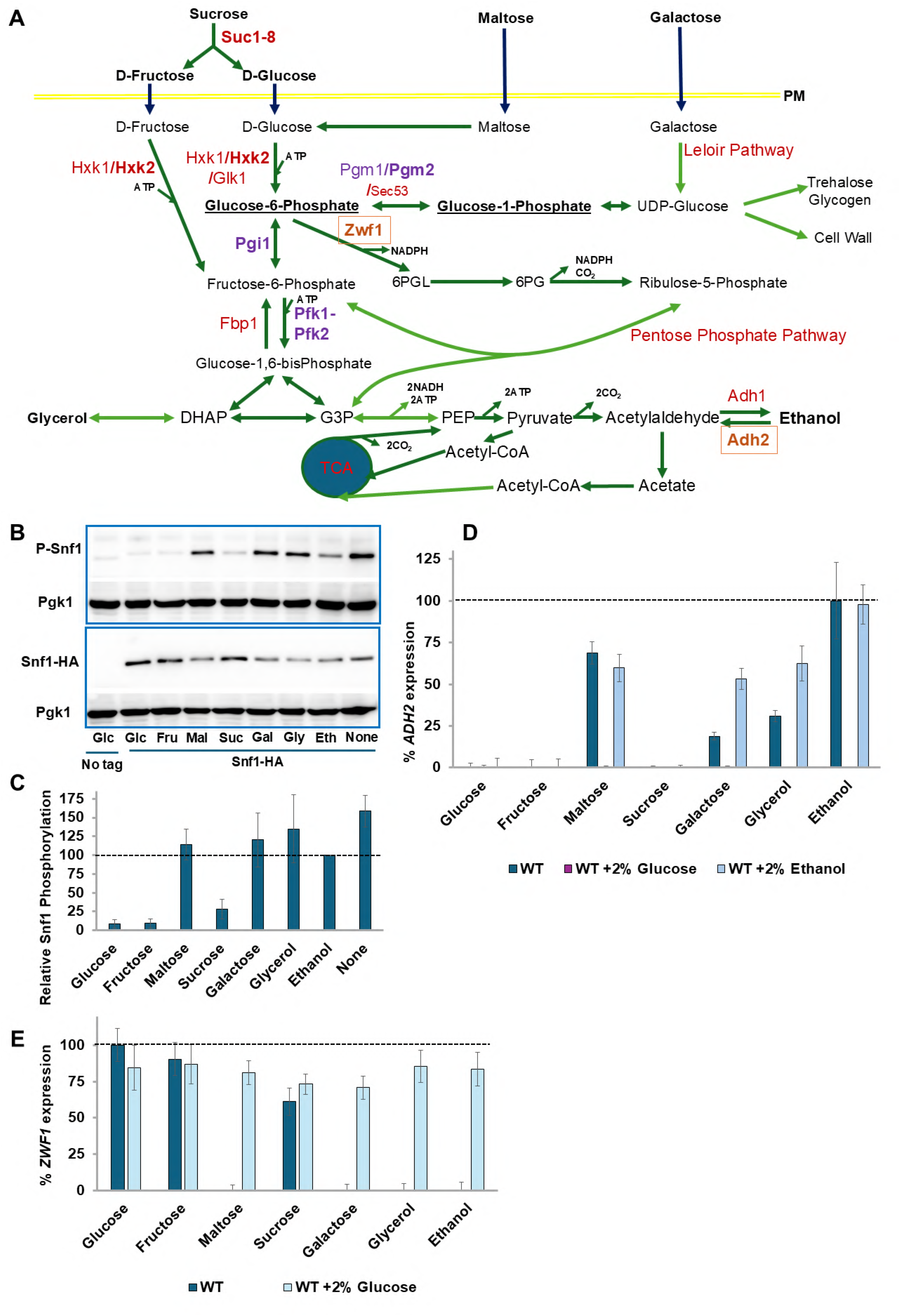
*ADH2* expression does not correlate with Snf1 activation. **A**. Schematic diagram illustrating carbon metabolic pathways of relevance to this article that are found in the V5 background. In **black bold** are carbon sources. **<u>Black Bold Underlined</u>** are Glucose-1-Phosphate (G1P) and Glucose-6-Phosphate (G6P). In **purple** are the enzymes whose expression is investigated in the article. In **red** are other important enzymes and Names of Metabolic Pathways. Dark Green arrows are single step reactions, whereas Light Green arrows indicate multiple steps (omitted for clarity). **A. B.** Glucose-grown cells were washed and then grown for 15 minutes in the indicated carbon source (**Glu**cose, **Fru**ctose, **Mal**tose, **Suc**rose, **Gal**actose, **Gly**cerol, **Eth**anol or **None**) as described in Materials and Methods. Cells were processed and Western Blots obtained as described in Materials and Methods. The experiment was repeated **four (maltose) or five** times. Pgk1 is used as a loading control. P-Snf1 is Snf1 phosphorylated at T210. **C**. Quantifications of Western Blots of **Figure 1B** were performed using ImageJ. **D.** Wild-type cells were grown overnight in 4% glucose, diluted and grown the following morning for an additional three hours. Basal *ADH2* expression was determined by β-galactosidase assay. Cells were washed and resuspended in media lacking any carbon source, and the indicated carbon sources added (also with an additional 2% Glucose or 2% Ethanol). *ADH2* expression was determined after three hours of growth at 30^◦^C. Expression rate is normalized to WT in ethanol. N=3**. E.** Wild-type cells were grown overnight in 4% glucose, diluted and grown the following morning for an additional three hours. Basal *ZWF1* expression was determined by β-galactosidase assay. Cells were washed and resuspended in media lacking any carbon source, and the indicated carbon sources added (also with an additional 2% Glucose or 2% Ethanol) with 0.05% MMS. *ZWF1* expression was determined after three hours of growth at 30^◦^C. Expression rate is normalized to WT in glucose. N=3.

### ZWF1 also shows differential regulation by carbon source

We also investigated other carbon source-regulated genes. *ZWF1* encodes *S. cerevisiae*’s G6PDH, which catalyzes the first step of the oxidative pentose-phosphate pathway (**Figure 1A**) and its expression is dramatically upregulated following DNA damage by methylmethane sulfonate (MMS) and other oxidative stresses [42]. *ZWF1* expression in response to MMS requires glucose or fructose (with a lower expression on sucrose) and is markedly repressed by maltose, galactose, glycerol and ethanol (**Figure 1D**). From a teleological perspective this is logical since the substrate of Zwf1 is Glucose-6-phosphate, so why produce this enzyme if the substrate is unavailable? However, this explanation does not provide a mechanism for how the different carbon sources are sensed. Adding 2% glucose to all poorer carbon source media restored *ZWF1* expression (**Figure 1D**) indicating that the inhibition of *ZWF1* expression is due to an **absence** of glucose (or fructose or sucrose) and not due to the **presence** of other carbon sources.

### Snf1, PKA and the SRR pathways do not regulate the carbon source differential *ADH2* and *ZWF1* expression levels

Snf1 activity is required for expression of *ADH2* by causing the dephosphorylation of the Adr1 transcription factor by an unidentified phosphatase. An Adr1^S230A^ mutant results in *ADH2* expression to occur in the absence of Snf1 [43]. However, expression of ADR1^S230A^ did not affect the expression of *ADH2* or *ZWF1* in the different carbon sources (**Figures S1D, S1E**). PKA is activated in *S. cerevisiae* by either Ras2 or Gpa2 sensing of glucose [44,45] . However, introduction of a constitutively active Ras2^G19V^ allele [44–46] does not affect *ADH2* or *ZWF1* expression on different carbon sources (**Figures S1D, S1E**). Another pathway that regulates genes that require glucose to be expressed is the SRR signaling pathway: glucose is sensed by Rgt2 and Snf3 relieving Rgt1 repression of (mainly hexose transporter) genes [4–6]. We examined the expression of *ADH2* (**Figure S1F**) and *ZWF1* (**Figure S1G**) in *rgt2Δ snf3Δ* cells. Inactivation of the SRR pathway does not affect the carbon source regulation of these genes.

Since there are no known signaling pathways that can discriminate between the different carbon sources, and indeed some carbon sources have no sensing mechanism (e.g. glycerol), we considered whether a metabolite could be responsible for these differences in gene expression, whereby the metabolite could potentially be a sensor for all carbon sources. Deletion of genes encoding specific glycolysis enzymes has previously been reported to cause accumulation of upstream metabolites when cells are grown on glucose (e.g. *TDH3* [47], *PFK2* [39]) or galactose (*PGM2* [39,48]).

### A balance of hexose metabolism regulates gene expression

An octameric complex of Pfk1 and Pfk2 [49] catalyzes the uni-directional glycolytic phosphorylation of fructose-6-phosphate by ATP to produce fructose-1-6-bisphosphate. The reverse reaction in gluconeogenesis is catalyzed by Fbp1 (**Figure S1A**). Single mutants of *PFK1 or PFK2* are viable on glucose (although with a lag phase during which they respire rather than ferment glucose [50] due to a throttling of glycolysis). However, both subunits are capable of catalyzing the phosphorylation of fructose-6-phosphate (F6P). [51]. We deleted *PFK1* since mutants of *PFK2* (but not of *PFK1*) also exhibit deficient Vma1 activity leading vacuolar alkalization and cytoplasmic acidification due to defective V-ATPase activity [52]. *pfk1Δ* cells exhibit a marked decrease in *ADH2* expression when maltose (a disaccharide made up of two glucose molecules), galactose or glycerol are the carbon source compared to wild-type cells (**Figure 2A**), suggesting that the accumulation of F6P or an upstream metabolite is inhibitory to *ADH2* expression. To distinguish between F6P and G6P, we used a *pgi1Δ* strain [53] and observed that a blockage of glycolysis at the interconversion of G6P to F6P also inhibits *ADH2* expression (**Figure 2B**) when cells are grown with maltose as the carbon source (which enters glycolysis as intracellular glucose); indicating that the metabolite in question is either G6P or one of its precursors. We therefore decided to restrict another exit point of G6P by preventing its reversible conversion to G1P by phosphoglucomutases (PGM).

**Figure 2.**
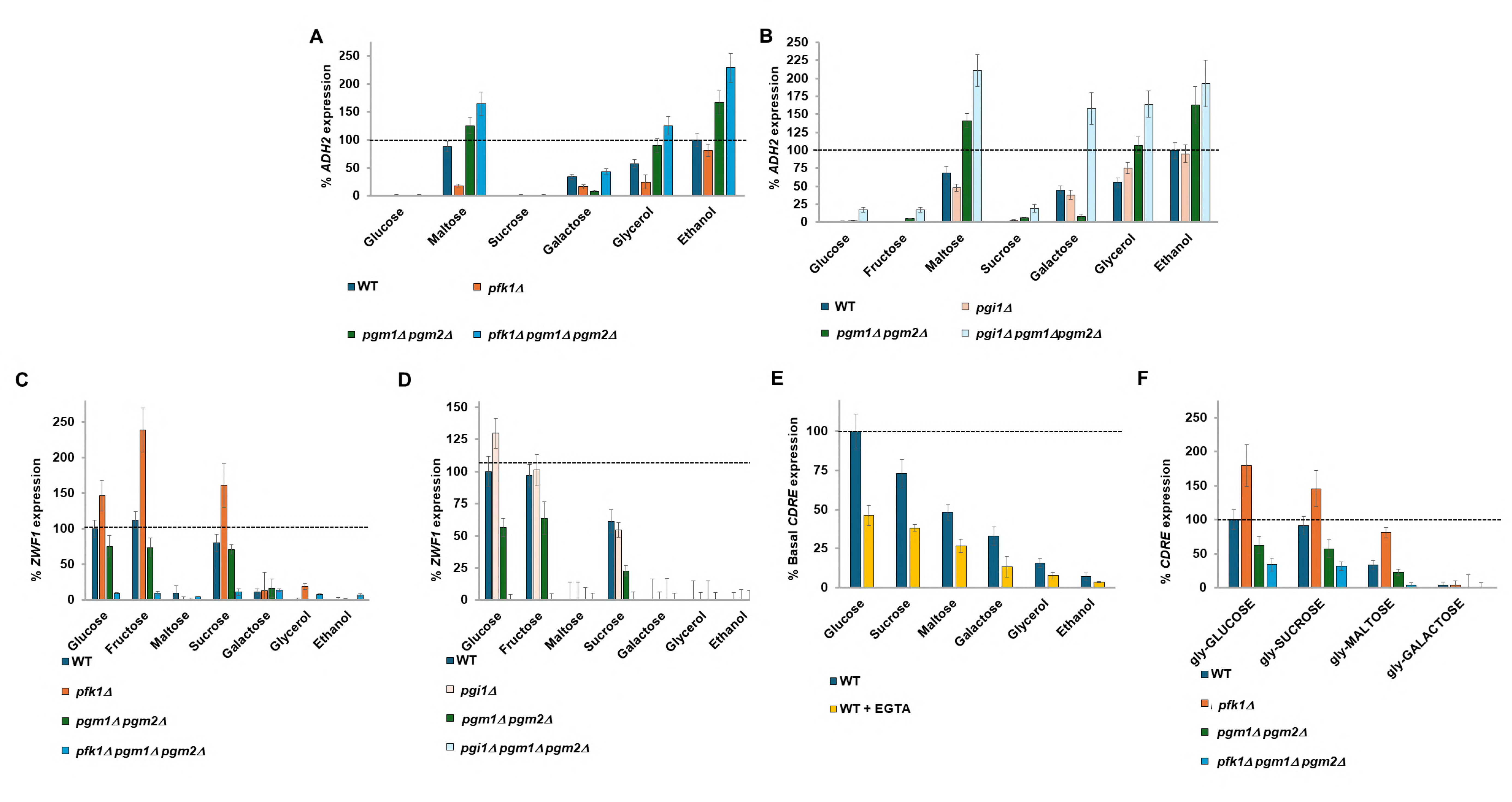
Deletions of glycolysis genes affect *ADH2, ZWF1* and *CDRE* expression. **A, B**. cells were grown overnight in 4% glucose (**A**) or 2% Fructose + 0.05% glucose (**B**), diluted and grown the following morning for an additional three hours. Basal *ZWF1* expression was determined by β-galactosidase assay. Cells were washed and resuspended in media lacking any carbon source, and the indicated carbon sources added (also with an additional 2% Glucose or 2% Ethanol) with 0.05% MMS. *ZWF1* expression was determined after three hours of growth at 30^◦^C. Expression rate is normalized to WT in glucose. N=3. cells were grown overnight in 4% glucose, diluted and grown the following morning for an additional three hours. Basal *ADH2* expression was determined by β-galactosidase assay. Cells were washed and resuspended in media lacking any carbon source, and the indicated carbon sources added (also with an additional 2% Glucose or 2% Ethanol). *ADH2* expression was determined after three hours of growth at 30^◦^C. Expression rate is normalized to WT in ethanol. N=3**. C, D.** cells were grown overnight in 4% glucose (**C**) or 2% Fructose + 0.05% glucose (**D**), diluted and grown the following morning for an additional three hours. Basal *ZWF1* expression was determined by β-galactosidase assay. Cells were washed and resuspended in media lacking any carbon source, and the indicated carbon sources added with 0.05% MMS. *ZWF1* expression was determined after three hours of growth at 30^◦^C. Expression rate is normalized to WT in glucose. N=3**. E**. Wild-type yeast were grown overnight at 30°C in selective media containing 2% of the carbon source indicated (glycerol at 3%), with or without the calcium chelator EGTA (1mM). Cells were diluted in the morning into the same carbon sources (and more EGTA added where required) and grown for an additional 3 hours before the basal *CDRE* expression was determined by β-galactosidase assay. Expression is normalized to WT in glucose. N=3. **F**. cells were grown overnight at 30°C in selective media containing 3% glycerol + 0.1% glucose. Cells were aliquoted into tubes and diluted in the morning into media containing 3% glycerol (no glucose) and grown for 3 hours. Samples were taken for determining *CDRE* expression under basal conditions (t=0). Indicated carbon sources (UPPERCASE) were added to 2% and the cells grown for 90 minutes before *CDRE* expression was determined by β-galactosidase assay (t=1.5). Expression is normalized to WT in glucose. N=3.

The production of G1P is essential for viability, with G1P being converted to UDP-glucose, which is in turn used for cell wall construction or production of storage carbohydrates. Pgm2 is the major phosphoglucomutase in *S. cerevisiae* [48]. Despite G1P being essential for viability, *pgm1Δ pgm2Δ* cells are viable on glucose media due to a low level of PGM activity provided by Sec53 [54]. The Leloir pathway converts galactose to G1P which is converted to G6P by PGM to enter glycolysis (**Figure S1A**).

*pgm1Δpgm2Δ* cells show an increase in *ADH2* expression in maltose, glycerol and ethanol media presumably due to reduced production of G1P (by Sec53) (**Figure 2A, B**). However, *ADH2* expression in *pgm1Δ pgm2Δ* cells is diminished in galactose (**Figure 2A, B**) since galactose metabolism produces G1P. Removal of both metabolic exits of G6P in a *pgm1Δ pgm2Δ pfk1Δ* or a *pgm1Δ pgm2Δ pgi1Δ* triple mutant results in a big increase in *ADH2* expression (**Figures 2A, 2B**), suggesting that it is the balance of G6P and G1P that is regulating *ADH2* expression, with higher G6P promoting expression.

### *ZWF1* expression is also regulated by Glucose-6-phosphate and Glucose-1-phosphate

*ZWF1* expression following 0.05% MMS treatment requires the presence of glucose, fructose or sucrose as the carbon source, with low expression in maltose, galactose and other poor carbon sources (**Figures 2C, 2D**). The expression profile of *ZWF1* is the inverse of *ADH2*, with a *pfk1Δ* (**Figure 2C**) or a *pgi1Δ* (**Figure 2D**) strain exhibiting higher *ZWF1* expression, suggesting that a metabolite above G6P promotes *ZWF1* expression. It is noted that whereas fructose causes very high *ZWF1* expression in a *pfk1Δ* strain (in which fructose-6-phosphate catabolism is restricted), fructose does not increase *ZWF1* expression in a *pgi1Δ* strain (in which fructose-6-phosphate catabolism is unimpeded) (compare **Figures 2C** and **2D**). *pgm1Δ pgm2Δ* cells show lower *ZWF1* expression, suggesting that elevated G1P levels promote *ZWF1* expression, and blocking both phosphoglucomutases and either Pfk1 or Pgi1 activity reduces *ZWF1* expression to near zero (**Figures 2C, 2D**); suggesting that it is again the balance of G6P and G1P that is regulating *ZWF1* expression, with higher G1P promoting *ZWF1* expression, and higher G6P repressing *ZWF1* expression. Thus, we conclude that the ratio of G6P to G1P is responsible for both the positive and negative regulation of carbon source-affected genes.

### Calcium signaling is regulated by all carbon sources

The pattern of regulation of *ADH2,* and *ZWF1* expression by G1P and G6P levels bears a high resemblance to the regulation of calcium accumulation by these metabolites [39]. *FKS2* encodes a β-glucan synthase required for cell wall synthesis. Its promoter contains a 24bp calcium-dependent response element (CDRE) [55] which, when fused to a LacZ reporter, is used to determine Crz1 activity [27,55] and by extension nucleoplasmic calcium levels [27]. Basal *CDRE* expression was determined by growing cells in different carbon sources overnight and with dilution into fresh media containing the same carbon source for two hours. Basal *CDRE* expression is high in glucose and becomes progressively lower the further down the glycolysis the carbon source is (**Figure 2E**). *CDRE* expression is sensitive to 1mM EGTA (which chelates external calcium [27,56] showing that this *CDRE* expression is due to calcium. **This suggests that calcium signaling generates a different analogue response for every carbon source**. We took glycerol-grown cells and added either 2% of different carbon sources (Glucose, Sucrose, Maltose, Galactose) or the same carbon sources together with 2% glucose. Similarly to the basal expression experiment of **Figure 2E**, *CDRE* expression is dependent upon the carbon source added to glycerol grown cells (**Figure S2B**). Further addition of 2% glucose restores expression of *CDRE* when co-present with all other carbon sources, with the *CDRE* response to glucose furthermore showing dose-dependency (as glucose was added either to 2% or to 4%) (**Figure S2B**).

### Glucose metabolites signal to calcium

Different carbon sources were added to glycerol-grown cells and the increase in *CDRE* expression was measured after 90 minutes. *pfk1Δ* cells exhibit a high increase in *CDRE* expression upon addition of either 2% glucose, 2% sucrose or 2% maltose (but not 2% galactose), whereas *pgm1Δ pgm2Δ* cells exhibit about a one-third decrease of *CDRE* expression, with *pfk1Δ pgm1Δ pgm2Δ* cells showing an almost total abolishment of *CDRE* expression (**Figure 2F**). Treatment with the calcium chelator EGTA lowers *CDRE* expression by one-half (**Figure S2C**). Thus, the expression profile for *CDRE* resembles that of *ZWF1* and is opposite to that of *ADH2*, suggesting that this calcium release that is regulated by the G6P/G1P ratio is regulating expression (both positively and negatively) of *ZWF1* and *ADH2*

### Nucleo-cytoplasmic calcium levels both positively and negatively regulate carbon source-dependent gene expression

A number of calcium pumps regulate calcium availability in the cell. The Vcx1 and to a lesser extent Pmc1 calcium pumps remove calcium from the cytoplasm into the vacuole and are key for maintaining cytoplasmic calcium homeostasis [24]. Pmr1 pumps calcium into the Golgi [57] together with Gdt1 [58]. Cod1 pumps calcium into the endoplasmic reticulum [59].

We screened the five calcium pumps for effects on carbon source-dependent gene expression. Knockout of either *COD1, PMR1* or *VCX1* increases glucose-induced *CDRE* expression and this increase of expression is sensitive to the calcium chelator EGTA (1mM). Knockout of *GDT1*, *PMC1* or *YVC1* has no effect on glucose-induced calcium levels (**Figure 3A**). Knockout of *COD1, PMR1* or *VCX1* lowers *ADH2* expression (**Figure 3B**) and this is suppressed by prevention of calcium influx by addition of 10 mM MgCl_2_ [27](**Figure S3A**), showing that the phenotype is due to higher calcium levels in the nucleo-cytoplasm [27,60]. Knockout of *PMR1* or *VCX1* increases *ZWF1* (**Figure 3C**) expression, whereas knockout of *COD1* exhibits a milder increase in *ZWF1*with maltose as the carbon source (**Figure 3C**). These increases are also suppressed by 10 mM MgCl_2_ (**Figures S3A, S3B**). Furthermore, since deletion of **any** of these three calcium pumps affects gene expression, it is the nucleo-cytoplasmic calcium levels that are regulating the gene expression rather than the calcium levels of a particular organelle. These three same calcium pumps have previously been shown to affect cytoplasmic calcium levels and the TECC response to glucose. Deletion of *VCX1* decreases vacuolar storage of calcium and raises cytoplasmic calcium levels, whereas deletion of *COD1* or *PMR1* increases the TECC release of calcium in response to glucose. Similarly to our results, D’hooge et al. found that deletions of *GDT1* or *PMA1* have little effect on glucose-responsive calcium release [37]. Deletion of the vacuolar calcium release channel Yvc1 did not affect *CDRE* expression, as expected [27], and did not affect *ADH2* or *ZWF1* expression (**Figures 3A, B, C**).

**Figure 3.**
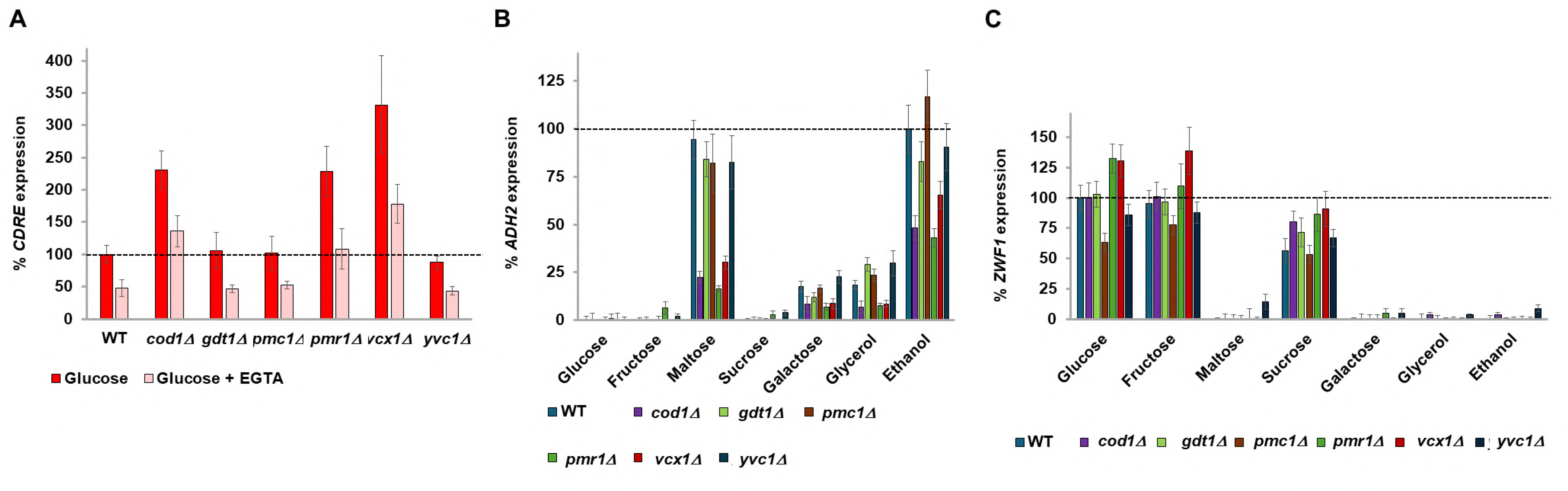
Deletions of calcium pumps regulate *CDRE, ADH2* and *ZWF1* expression. **A**. cells were grown overnight in media with 4% glucose (and 1mM EGTA where indicated). Cells were diluted in the morning with media with 4% glucose (and 1mM EGTA where indicated) and cells grown for an additional three hours before *CDRE* expression was determined by β-galactosidase assay. N=3. Expression is normalized to WT in glucose. **B**. cells were grown overnight in 4% glucose, diluted and grown the following morning for an additional three hours. Basal *ADH2* expression was determined by β-galactosidase assay. Cells were washed and resuspended in media lacking any carbon source, and the indicated carbon sources added. *ADH2* expression was determined after three hours of growth at 30^◦^C. Expression rate is normalized to WT in ethanol. N=3**. C.** cells were grown overnight in 4% glucose, diluted and grown the following morning for an additional three hours. Basal *ZWF1* expression was determined by β-galactosidase assay. Cells were washed and resuspended in media lacking any carbon source, and the indicated carbon sources with 0.05% MMS. *ZWF1* expression was determined after three hours of growth at 30^◦^C. Expression rate is normalized to WT in glucose. N=3.

### Raising nucleo-cytoplasmic calcium levels suppresses phenotypes caused by low G1P

We next decided to determine whether elevating nucleo-cytoplasmic calcium levels by knockout of *COD1* or *VCX1* would suppress the changes in gene expression observed in *pgm1Δ pgm2Δ* cells (*pgm1Δ pgm2Δ pmr1Δ* cells were not viable). Deletion of either *COD1* or *VCX1* restores expression of *CDRE* to at least WT levels (**Figure S4A**) suggesting that nucleo-cytoplasmic calcium levels are likewise restored in these cells. Deletion of either *COD1* or *VCX1* abolishes the increased *ADH2* expression of *pgm1Δ pgm2Δ* cells (**Figure S4B**), and suppresses the decreased *ZWF1* expression observed in *pgm1Δ pgm2Δ* cells (**Figure S4C**).

Since the phenotype of *pgm1Δ pgm2Δ* cells is sometimes rather mild (e.g. for *ZWF1* expression), we constructed *cod1Δ pfk1Δ pgm1Δ pgm2Δ* and *cod1Δ pgi1Δ pgm1Δ pgm2Δ* strains to determine whether raising nucleo-cytoplasmic calcium levels could suppress the severe phenotypes of blocking both metabolic exits of Glucose-6-phosphate (**Figure S1A**). Further deletion of *COD1* restores nucleo-cytoplasmic levels of *CDRE* expression of *pfk1Δ pgm1Δ pgm2Δ* cells in glucose, sucrose and maltose media to WT levels (**Figure 4A**). Further deletion of *COD1* likewise suppresses the increased *ADH2* (**Figure 4B**), and decreased *ZWF1* (**Figure 4C**) expression of *pfk1Δ pgm1Δ pgm2Δ* cells, with similar results for *ADH2* (**Figure S4D**) and *ZWF1* (**Figure S4E**) obtained when *PGI1* was deleted in lieu of *PFK1.* Although deletion of *COD1* alone does not drastically affect *ZWF1* expression (**Figures 3C, 4C, S4C**), deletion of *COD1* suppresses the low *ZWF1* expression phenotype of *pgm1Δ pgm2Δ* and of *pfk1Δ pgm1Δpgm2Δ* cells. Jointly, our results demonstrate that increased nucleo-cytoplasmic calcium levels suppress the phenotypes of dysregulated G1P and G6P levels.

**Figure 4.**
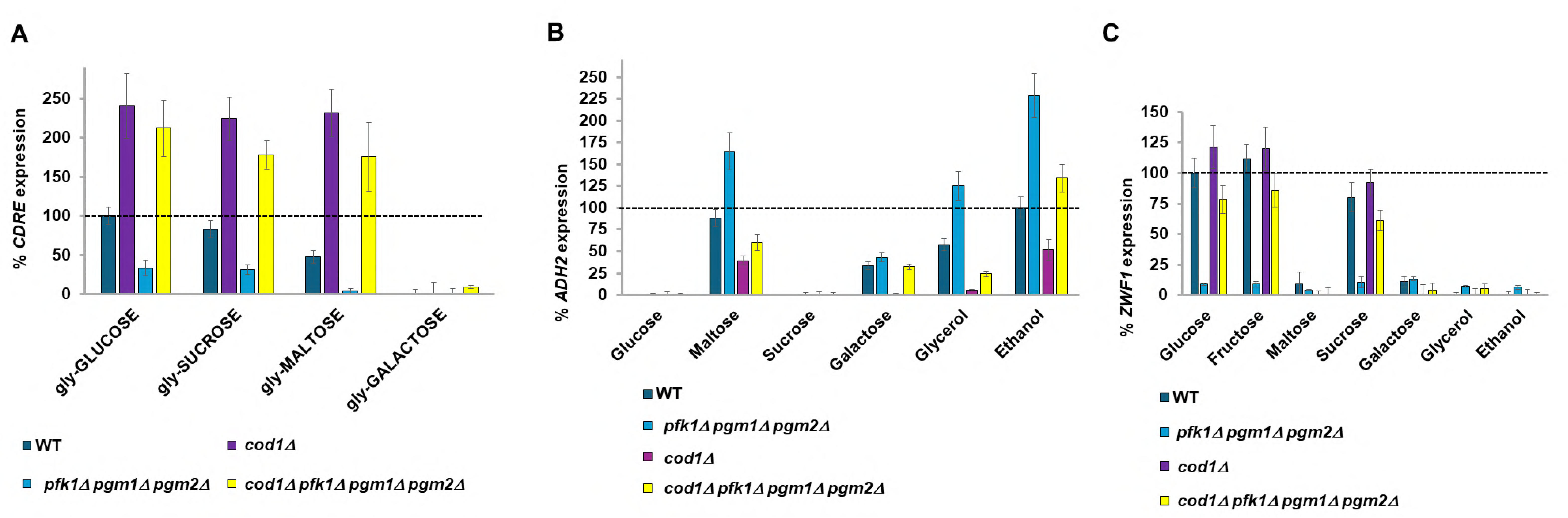
Deletion of *COD1* or *VCX1* suppresses phenotypes of *pfk1Δpgm1Δpgm2Δ* cells. **A**. cells were grown overnight at 30°C in selective media containing 3% glycerol + 0.1% glucose. Cells were aliquoted into tubes and diluted in the morning into media containing 3% glycerol (no glucose) and grown for 3 hours. Samples were taken for determining *CDRE* expression under basal conditions (t=0). Indicated carbon sources (UPPERCASE) were added to 2% and the cells grown for 90 minutes before *CDRE* expression was determined by β-galactosidase assay (t=1.5). Expression is normalized to WT in glucose. N=3. **B**. cells were grown overnight in 4% glucose, diluted and grown the following morning for an additional three hours. Basal *ADH2* expression was determined by β-galactosidase assay. Cells were washed and resuspended in media lacking any carbon source, and the indicated carbon sources added. *ADH2* expression was determined after three hours of growth at 30^◦^C. Expression rate is normalized to WT in ethanol. N=3**. C.** cells were grown overnight in 4% glucose, diluted and grown the following morning for an additional three hours. Basal *ZWF1* expression was determined by β-galactosidase assay. Cells were washed and resuspended in media lacking any carbon source, and the indicated carbon sources with 0.05% MMS. *ZWF1* expression was determined after three hours of growth at 30^◦^C. Expression rate is normalized to WT in glucose. N=3.

### The G1P/G6P ratio signals to Pma1 proton pumping

Addition of glucose to media not only results in transient calcium influx into the nucleo-cytoplasm, but also activates the Pma1 proton pump [61], with Pma1 activity being required for calcium signaling but not vice-versa [61]. Surprisingly, the mechanisms by which both Pma1 and the TECC response are activated following glucose addition are unknown.

In a previous study, we genetically manipulated Pma1 activity using a non-inhibitable Pma1-*Δ*901 allele coupled with deletion of *HSP30* to elevate Pma1 activity [62] and an *EXP1* deletion to lower the amount of Pma1 at the plasma membrane. Here we continue to utilize *PMA1-Δ901 hsp30Δ* but have also introduced the *hrk1Δ ptk2Δ* double knockout which cannot activate Pma1 in response to glucose [7]. We also utilize the *psg1Δ* knockout which exhibits lower levels of plasma membrane-localized Pma1 [63].

Inhibition of Pma1 activity by deletion of *HRK1* and *PTK2* [7] decreases *CDRE* expression (calcium release) when glycerol-grown WT cells are fed glucose, sucrose or maltose and lowers the high *CDRE* expression of *pfk1Δ* cells (**Figure 5A**). Lowering Pma1 levels at the plasma membrane by deletion of *PSG1* [63] likewise lowers *CDRE* expression. Inhibition of Pma1 by these methods increases *ADH2* expression (**Figure 5B**) and decreases *ZWF1* expression (**Figure 5C**). Moreover, it also suppresses the decreased *ADH2* (**Figure 5B**) and increased *ZWF1* expression (**Figure 5C**) of *pfk1Δ* cells. Hyper-activation of Pma1 (*PMA1-Δ901 hsp30Δ*) dramatically increases *CDRE* expression upon addition of glucose, sucrose and even maltose (**Figure S5A**) and suppresses the low *CDRE* expression phenotype of *pgm1Δ pgm2Δ* cells (**Figure S5A**)

**Figure 5.**
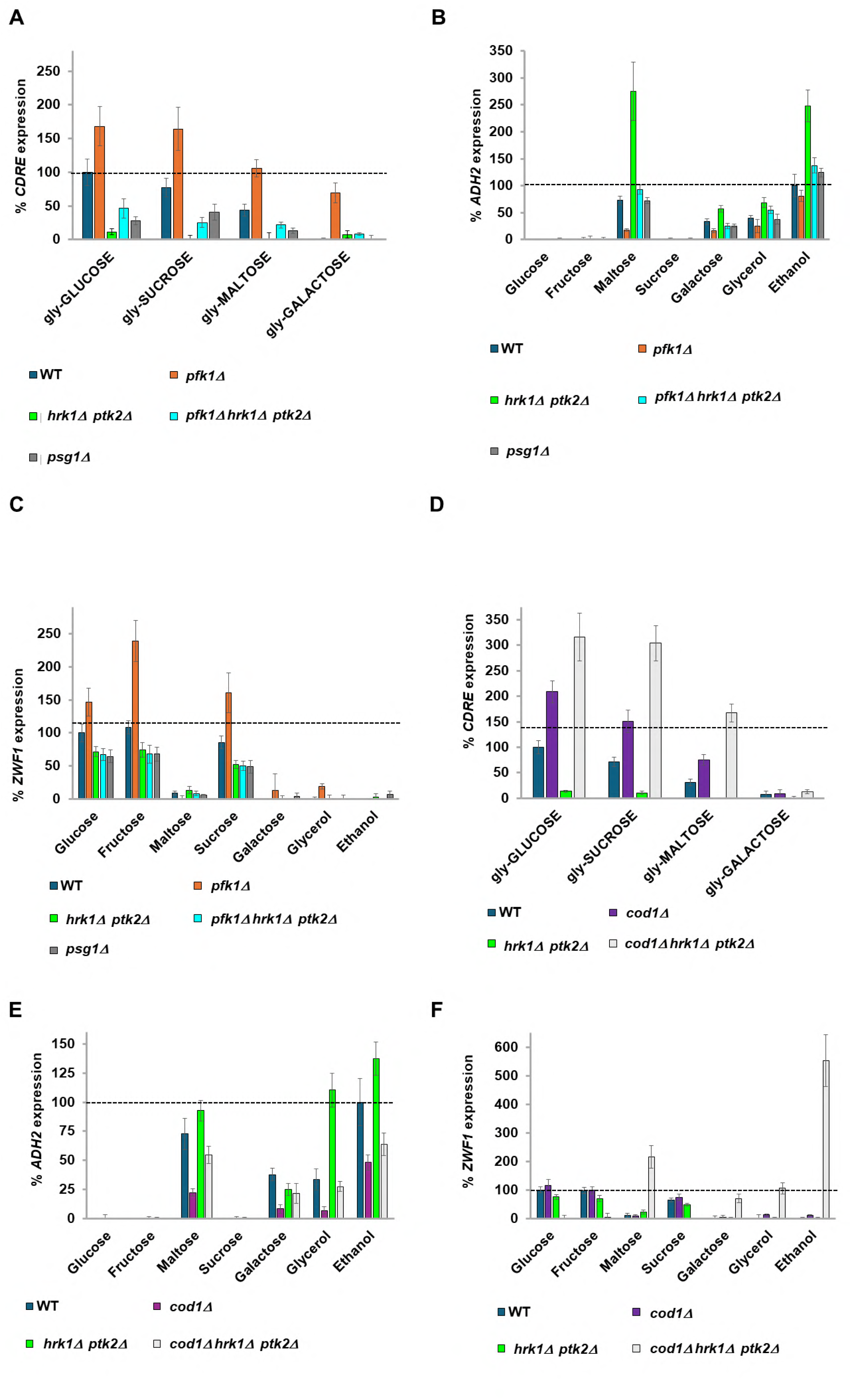
Manipulation of Pma1 activity suppresses *pfk1Δ* phenotypes. **A, D**. cells were grown overnight at 30°C in selective media containing 3% glycerol + 0.1% glucose. Cells were aliquoted into tubes and diluted in the morning into medium containing 3% glycerol (no glucose) and grown for 3 hours. Samples were taken for determining *CDRE* expression under basal conditions (t=0). Indicated carbon sources (UPPERCASE) were added to 2% and the cells grown for 90 minutes before *CDRE* expression was determined by β-galactosidase assay (t=1.5). Expression is normalized to WT in glucose. N=3. **B, E**. cells were grown overnight in 4% glucose, diluted and grown the following morning for an additional three hours. Basal *ADH2* expression was determined by β-galactosidase assay. Cells were washed and resuspended in media lacking any carbon source, and the indicated carbon sources added. *ADH2* expression was determined after three hours of growth at 30^◦^C. Expression rate is normalized to WT in ethanol. N=3**. C, F.** cells were grown overnight in 4% glucose, diluted and grown the following morning for an additional three hours. Basal *ZWF1* expression was determined by β-galactosidase assay. Cells were washed and resuspended in media lacking any carbon source, and the indicated carbon sources with 0.05% MMS. *ZWF1* expression was determined after three hours of growth at 30^◦^C. Expression rate is normalized to WT in glucose. N=3.

Pma1 hyper-activation (*PMA1-Δ901 hsp30Δ*) decreases *ADH2* expression and suppresses the high *ADH2* expression of *pgm1Δ pgm2Δ* cells (**Figure S5B)**, indicating that G6P/G1P sensing precedes Pma1 for the regulation of *ADH2.* In contrast, reduction of Pma1 activity whether by prevention of Pma1-tail phosphorylation (*hrk1Δ ptk2Δ*) or by decreasing its trafficking to the plasma membrane (*psg1Δ*) increases *ADH2* expression (**Figure S5B**), although *psg1Δ* has a more limited effect than prevention of Pma1-tail phosphorylation. Similarly to deletion of *COD1* (**Figures 3C, 4C, S4C**), **h**yper-activation of Pma1 (*hsp30Δ PMA1-Δ901*) alone does not affect *ZWF1* expression but suppresses the low *ZWF1* expression of *pgm1Δ pgm2Δ* cells (**Figure S5C**). Together these results suggest two possible models. The first is that the G6P/G1P ratio signaling is upstream of, and regulates, Pma1 activity. The second possibility is that the forced Pma1-induced changes to cytoplasmic pH overwhelm the metabolite-generated signal.

To differentiate between these possibilities, we determined whether upregulating calcium signaling could compensate for a lack of Pma1 activity. Additional deletion of the *COD1* calcium pump totally suppresses the *CDRE* expression defect of *hrk1Δptk2Δ* cells (**Figure 5D**). Deletion of *COD1* also lowers the *ADH2* hyper-expression phenotype of *hrk1Δ ptk2Δ* cells, though not to the same level as deletion of *COD1* in cells with functional Hrk1 and Ptk2 (**Figure 5E**). For *ZWF1* expression, a *hrk1Δ ptk2Δ cod1Δ* strain exhibits an inverted expression profile with low expression in glucose and fructose media and yet very high expression in maltose, galactose, glycerol and ethanol media (**Figure 5F**). Taken together, these results show that the phenotypes observed are due to calcium dysregulation in the absence of Pma1 activity.

### Signaling downstream of calcium bifurcates

Research into signaling downstream of calcium has been extremely biased towards the calcineurin phosphatase and its substrate Crz1, which it activates. Calcineurin inhibits Vcx1 pump activity [64] but Crz1 upregulates *PMC1* and *PMR1* transcription [55]. Other targets of calcium include the calmodulin-activated kinases (Cmk1, Cmk2 and Rck1). Treatment of glycerol-grown WT cells with 1mM of the calcineurin inhibitor FK506 [55] just before addition of glucose, sucrose or maltose drastically lowers *CDRE* expression, as expected (**Figure 6A**). Deletion of *CMK1* has no effect on *CDRE* expression, as expected (**Figure 6A**). Administration of 1mM FK506 immediately after washing cells before resuspension in different carbon sources increases *ADH2* expression, with cells on media with maltose, galactose and glycerol now having the same expression level as those in ethanol (**Figure 6B**). This suggests that in the absence of calcineurin signaling, the increased calcium levels (inferred by *CDRE* expression) of maltose-, galactose- and glycerol-(compared to ethanol-) grown cells (**Figure 2E**) no longer affect *ADH2* expression. Deletion of *CMK1* has no effect on *ADH2* expression (**Figure 6B**), but shows a marked decrease in *ZWF1* expression (**Figure 6C**). Thus, the downstream effector that regulates *ADH2* expression is calcineurin, whereas the downstream effector that regulates *ZWF1* is Cmk1.

**Figure 6.**
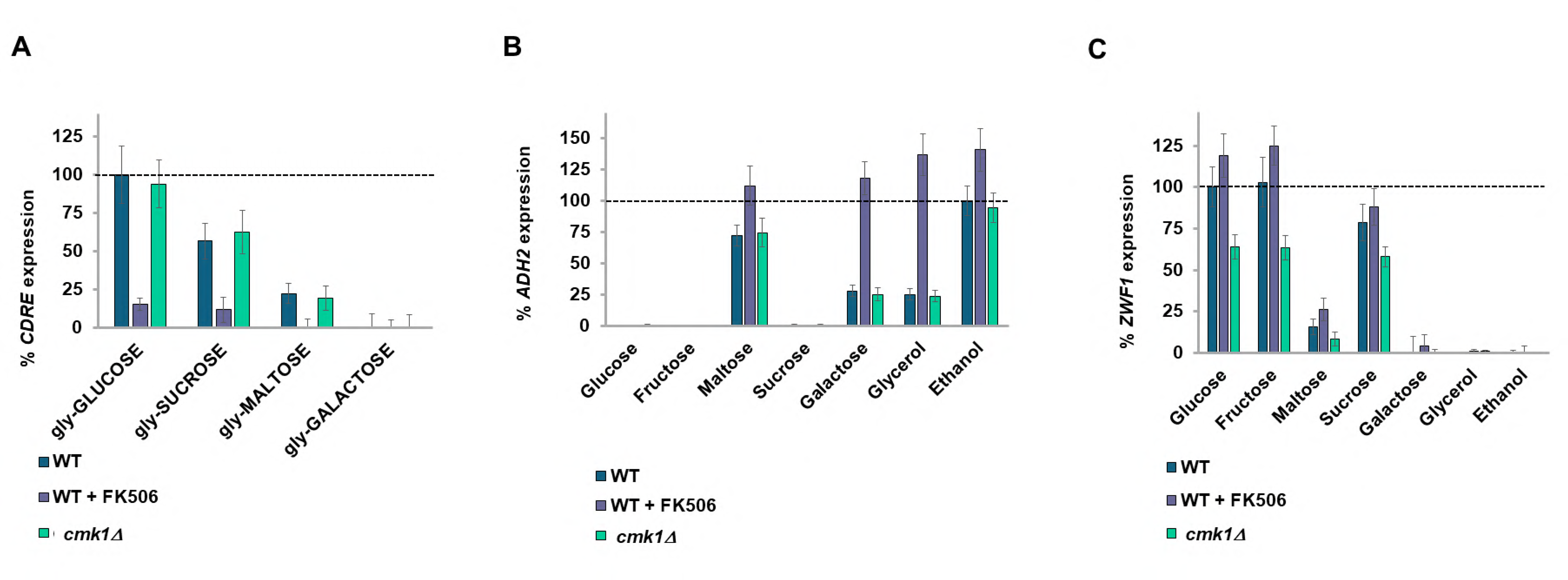
Signaling downstream of calcium is bifurcated. **A**. cells were grown overnight at 30°C in selective media containing 3% glycerol + 0.1% glucose. Cells were aliquoted into tubes and diluted in the morning into media containing 3% glycerol (no glucose) and grown for 3 hours. Samples were taken for determining *CDRE* expression under basal conditions (t=0). Indicated carbon sources (UPPERCASE) were added to 2% and 1mM FK506 were indicated and the cells grown for 90 minutes before *CDRE* expression was determined by β-galactosidase assay (t=1.5). Expression is normalized to WT in glucose. N=3. **B**. cells were grown overnight in 4% glucose, diluted and grown the following morning for an additional three hours. Basal *ADH2* expression was determined by β-galactosidase assay. Cells were washed and resuspended in media lacking any carbon source, and the indicated carbon sources added 1mM FK506 where indicated. *ADH2* expression was determined after three hours of growth at 30^◦^C. Expression rate is normalized to WT in ethanol. N=3**. C.** cells were grown overnight in 4% glucose, diluted and grown the following morning for an additional three hours. Basal *ZWF1* expression was determined by β-galactosidase assay. Cells were washed and resuspended in media lacking any carbon source, and the indicated carbon sources with 0.05% MMS and 1mM FK506 where indicated. *ZWF1* expression was determined after three hours of growth at 30^◦^C. Expression rate is normalized to WT in glucose. N=3.

### Calcium storage provides a memory mechanism of previous meals

Growth in different carbon sources results in a differential level of basal cytoplasmic calcium (inferred by *CDRE* expression) in WT V5 cells (**Figure 2E**) as well as in WT BY4741 cells (**Figure S6B**). The basal *CDRE* expression is decreased in *pgm1Δ pgm1Δ* (and even more so in *pfk1Δ pgm1Δ pgm2Δ*) cells. Deletion of *COD1* dramatically elevates basal *CDRE* expression and suppresses the low basal *CDRE* expression of *pfk1Δ pgm1Δ pgm2Δ* cells (**Figure 7A**). As expected, the effects of *COD1* deletion are strongest when cells are grown with a carbon source that produces a lot of glucose-phosphates (V5 cells have a growth defect on galactose, **Figure S6A**). We now wondered whether the calcium response to glucose would be the same irrespective of which carbon source the cells had been previously grown in. Upon addition of 2% glucose, cells previously grown in glucose or sucrose exhibit the highest increase in *CDRE* expression (calcium release), with cells prior-grown on galactose having 50% and other glycerol/ethanol prior-grown cells 20% of the *CDRE* expression of glucose prior-grown cells (**Figure 7A**). *pfk1Δ, pgm1Δ pgm2Δ* and *pfk1Δ pgm1Δ pgm2Δ* cells all exhibit differential *CDRE* expression upon addition of2% glucose depending upon the previous carbon source although the amplitude of *CDRE* expression is greater than wild-type for *pfk1Δ* cells across all prior carbon sources upon addition of 2% glucose and is lower than wild-type for *pgm1Δ pgm1Δ* and *pfk1Δ pgm1Δ pgm2Δ* cells upon addition of 2% glucose **(Figure 7B**). Deletion of *COD1* strongly elevates *CDRE* expression upon addition of 2% glucose across all prior carbon sources and suppresses the low *CDRE* expression phenotype of *pfk1Δ pgm1Δ pgm2Δ* cells (**Figure 7B**). Remarkably, the degree of *CDRE* induction upon addition of 2% glucose correlates strongly with the basal CDRE expression levels.

**Figure 7.**
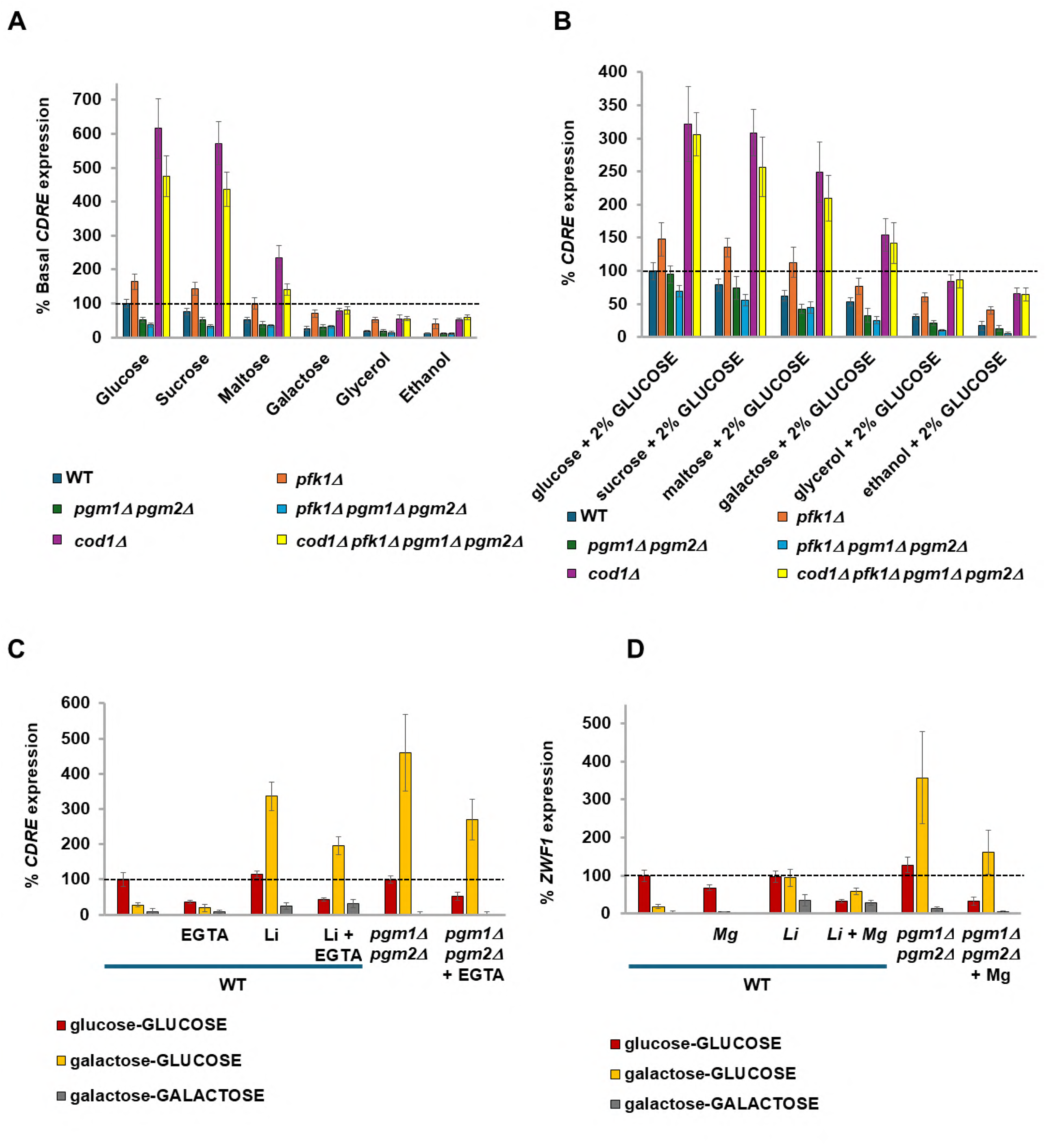
Calcium signaling serves as an ionic memory of prior meals. **A.** Cells were grown in the indicated carbon source (2%, except glycerol at 3%) diluted with media containing the same carbon source and grown for 3 hours before *CDRE* expression was determined by β-galactosidase assay. Expression is normalized to WT in glucose. N=3. **B**. 2% glucose was added to the cells from **Figure 7A** and *CDRE* expression determined by β-galactosidase assay after an additional 90 minutes. Expression is normalized to WT in glucose. Note that the *CDRE* expression of the glycerol-GLUCOSE condition is the normalizing condition for previous *CDRE* experiments. N=3. **Figures 7B, C**. BY4741 cells expressing β−galactosidase from either a *CDRE* (**Figure 7B**) or a *ZWF1* (**Figure 7C**) promoter were grown overnight at 30°C in selective media containing 3% glycerol + 0.1% glucose. Cells were diluted 5-fold and grown for 6 hours in media containing 3% glycerol (no glucose). WT cells to be treated with 15mM LiCl and to be grown in media containing 2% galactose or *pgm1Δpgm2Δ* cells to be grown in media containing 2% galactose were diluted 4x. Other cells were diluted 50x. Cells were distributed into tubes and grown under indicated conditions (combinations of 2% glucose, 2% galactose, 15mM LiCl, 1mM EGTA (**Figure 7B**)) for 16 hours and the basal β-galactosidase activity measured (t=0). For **Figure 7C** (*ZWF1* expression), 0.05% MMS was now added. The glucose grown cells now had another 2% glucose added to them (glucose-GLUCOSE), the galactose cells now had either 2% glucose (galactose-GLUCOSE) or 2% galactose (galactose-GALACTOSE) added. 10mM MgCl2 was added as indicated (**Figure 7C**). After 90 minutes (**Figure 7B**) or 3 hours (**Figure 7C**) the β-galactosidase activity was determined. Note that all the WT cells used in each experiment were from the same colony grown in 3% glycerol + 0.1% glucose. Expression rates are normalized to WT glucose-GLUCOSE (**Figure 7B**) or to WT glucose (**Figure 7C**). N=3

Since the V5 background has a growth defect on galactose and yet V5 *pgm1Δ pgm2Δ* cells are viable on galactose media (**Figure S6A**), we utilized the BY4741 background which grows well on galactose media. We took the cells from **Figure S6B** which had been grown overnight in different carbon sources and determined the *CDRE* expression after addition of 2% glucose. Similarly to V5 cells (**Figure 7A**), expression of *CDRE* in BY4741 cells is also dependent upon the prior carbon source, and the increases of *CDRE* expression are sensitive to EGTA (**Figure S6C**). Since the “basal” cells have had two exposures to their overnight carbon source (once upon seeding of cells, and again when diluted in the morning and grown for 3 hours) this suggests that the exposures to carbon sources (especially glucose) has resulted in post -TECC pumping of calcium into the stores [37], and thus prior exposure to glucose has led to an enhanced calcium release (as occurred in **Figures 7B, S6C**), with calcium serving as an **ionic memory** of past carbon source exposures. This concept is now extended to all carbon sources, with the intensity of the response diminishing the further metabolically the carbon source is from glucose-phosphates.

To show that increased calcium pumping due to a previous meal increases the intracellular levels of calcium and thus the TECC response, we decided to manipulate the cells to accumulate calcium by using the calcium hyperaccumulation phenotype of galactose-grown cells either deleted of *PGM* [39,48] or with PGM inhibited with 15 mM Lithium Chloride [40,65] .

Since calcium accumulation in *pgm1Δpgm2Δ* cells or WT cells treated with 15 mM LiCl is a very slow process [65], cells were grown in glycerol before being grown in either glucose or galactose media overnight in the presence of 15 mM lithium (and 1 mM EGTA) where indicated, before addition of an additional 2% glucose or galactose as indicated for determination of the increase in *CDRE* expression.

Lithium or knockout of *pgm1Δ pgm2Δ* did not affect *CDRE* expression upon addition of glucose to cells prior-grown in glucose, as would be expected since there is no increase in galactose-1-phosphate or calcium hyper-pumping. Addition of galactose to galactose prior-grown cells did not result in *CDRE* expression. However, lithium treated WT cells or *pgm1Δ pgm2Δ* cells pre-grown in galactose have an approximately 10-15 fold increase of *CDRE* expression upon treatment with 2% glucose compared to cells not treated with lithium or possessing *PGM1 PGM2* (**Figure 7C**). Thus, we have divorced calcium accumulation from growth in glucose, using the calcium hyper-pump to create a false memory of prior food.

Does this apply to other genes? We adapted this protocol for examining *ZWF1* expression. Similarly to *CDRE* expression, we found that overnight growth of lithium treated or *pgm1Δ pgm2Δ* cells in galactose results in high *ZWF1* expression upon release of calcium by addition of 2% glucose to the media (**Figure 7D**). Thus, this memory mechanism also regulates other calcium regulated genes.

## Discussion

We have uncovered a molecular mechanism that allows cells to adapt their metabolic gene expression to match any carbon source. This system converts the presence of each carbon source into an analogue calcium output (as best visualized in **Figures 2E** and **S6B**). This mechanism does not override glucose-based catabolite repression, but within the set of permissive carbon sources, it regulates the degree of gene expression. For *ADH2* expression this translates into Snf1 being active [43], and then calcineurin activity determines the precise level of expression, by analogy to a radio whereby firstly it must be switched on (by Snf1) and then the volume adjusted (by calcineurin). Hyperactivation of Adr1 by using an Adr1^S230A^ allele [43] does not affect the permissive carbon source regulation of *ADH2*. Inhibition of *ADH2* expression by increasing cytoplasmic concentrations of calcium, whether by metabolic (deletion of *PFK1* or *PGI1*) or calcium pump manipulation (deletion of *COD1*) indicates a role for calcium in the regulation of *ADH2* expression, with calcineurin activity being sufficient in galactose- and glycerol-grown cells to lower *ADH2* expression as compared with cells grown with ethanol as the carbon source. Although Crz1 is the best-characterized transcription factor working downstream from calcineurin, the *ADH2* promoter does not have a Crz1 binding site, suggesting that calcineurin communicates with other transcription factors. For *ZWF1* expression, Rpn4 is activated by MMS (or Yap1 by oxidative stress) [42] and then the extent of *ZWF1* expression is determined by Cmk1 activity.

We also found that this mechanism not only regulates glucose-repressed gene expression, but also glucose-dependent gene expression, such as *ZWF1* induction following DNA damage. From a teleological perspective it is logical for *ZWF1* to be expressed only when its substrate (Glucose-6-Phosphate) is present in sufficient quantity, and this mechanism of conversion of a G1P signal (which in wild-type cells *is* freely convertible with G6P) to a calcium level allows yeast to detect such carbon sources even in the absence of specific sensor. Increasing cytoplasmic calcium levels whether by metabolic or calcium pump manipulations increases *ZWF1* expression. Calcium activates Cmk1, whose activity increases *ZWF1* expression above the basal sucrose level when cells are grown with glucose or fructose as the carbon source. In addition, when Pma1 phosphorylation (and activation) was prevented by deletion of *HRK1* and *PTK2* **and** cytoplasmic calcium levels raised by deletion of *COD1* (**Figure 5F**) or *VCX1* (data not shown), an inverted phenotype occurred with repression of *ZWF1* expression on normally permissive carbon sources (glucose, fructose, sucrose) and very high expression on normally non-permissive carbon sources (especially ethanol). Other genes (*CLN2, BCK2, CDC28)* have previously been found to be induced by glucose upon refeeding and are also regulated by accumulation of glucose-phosphates, and similarly to *ZWF1* are not regulated by the SRR pathway [66] – perhaps these genes are also calcium regulated.

Most research into the calcium response to carbon sources has hitherto focused on glucose (e.g. [27]), although galactose has also been examined, usually in the context of Pgm2 inhibition [19]. In our hands, although glucose and sucrose addition to glycerol-grown cells resulted in high calcium signaling and permitted *ZWF1* expression whilst inhibiting *ADH2* expression, galactose resulted in a much-reduced calcium signal and did not permit *ZWF1* expression or totally repress *ADH2* expression.

An advantage of using a β-galactosidase reporter system is that the long half-life of the β-galactosidase protein results in a system that is sensitive to small changes. By growing cells overnight in different carbon sources, we were thus able to observe even small differences in the calcium response between galactose-, glycerol- and ethanol-grown cells (**Figures 2E, S6B**) which are presumably unobservable by TECC measurements or GFP based reporters. This demonstrates that **all** carbon sources have a demonstrable effect on basal levels of cytoplasmic calcium, thus broadening the scope of calcium signaling. The mechanism by which glycolysis metabolite levels transduce to a calcium signal is not known. Previous work revealed that it is the balance of G1P (pro-calcium) and G6P (anti-calcium) that regulates TECC [39] and presumably calcineurin activity [27] This work expands upon this model to include potentially all carbon sources, with those furthest from PGM activity exerting the least influence. This mechanism also provides a sensing mechanism for those carbon sources without specific sensors, such as sucrose, glycerol or ethanol. The extent of gene regulation by carbon source via calcium is also expanded beyond the traditionally investigated Crz1 targets to include both positive and negatively affected genes (*ZWF1* and *ADH2*). The full gamut of genes regulated by calcium in response to carbon source is likely to be much higher, since 32 carbohydrate metabolism genes are regulated by Cnb1 and Crz1 in response to media alkalization stress [67]. We also found comparable results to *ZWF1* expression for *HXT3* expression (data not shown).

Furthermore, the response to 2% glucose is not uniform, but rather depends on the carbon source the yeast is growing in (**Figure 7B, S6B**). This can arise since the pumping of calcium ions out of the cytoplasm is itself stimulated by glucose [37] and thus successive exposures to glucose would result in a feed-forward effect with ever-increasing calcium pumping and release upon the next glucose exposure. This differential release of calcium upon exposure to glucose dependent upon the previous carbon source could be considered an **ionic memory** mechanism whereby future responses to carbon source are dependent upon past meals (**Figure 7C, D**). Other examples of protein-based memory systems for carbon sources include the sequestration of Std1 into puncta in glucose-grown cells resulting in an increased *ADH2* expression when cells are transferred to glycerol compared with galactose grown cells [68], Whi5 protein determining the time of G1 depending upon the environmental conditions of previous G1 stages [69] or the accelerated expression of *GAL* genes upon transfer of cells from glucose to galactose **if** the cells had been previously exposed to galactose in the previous twelve hours [70].

Following addition of glucose to cells, Pma1 is rapidly disinhibited by phosphorylation on the C-terminal inhibitory tail by the Ptk2 and Hrk1 kinases [7] [upon withdrawal of glucose these sites are dephosphorylated by Glc7-Reg1 [71]]. The mechanism by which glucose activates these kinases is not known. Pma1 activation is required for TECC response to glucose [61]. Therefore, we sought to integrate Pma1 activity into our schema of Glucose-1-Phosphate/Glucose-6-Phosphate somehow activating calcium signaling. We found that Pma1 hyper-activation (achieved by deleting *HSP30* and using a tail-less Pma1-*Δ*901 mutant [62]) suppresses the effects of altering Glucose-6-Phosphate/Glucose-1-Phosphate levels by the glycolysis mutants on *CDRE, ADH2* and *ZWF1* expression (**Figures 5A-C**, **S5A-C**), and that deletion of *COD1* in turn suppresses the effects of Pma1 inhibition (**Figures 5D-F**). This concords with previous work [61], but it is not clear from these data whether the G6P/G1P ratio is involved in Pma1 activation or whether Pma1 activity is simply more significant a regulator of calcium release. At each of these stages of metabolite balance, Pma1 activation and calcium release the molecular mechanism remains to be determined.

In mammalian cells, glucose stimulates calcium influx in brain tissue (glia, neurons, astrocytes) [72] and in the pancreas [73]; whereas lactate (a poor carbon source that enters glycolysis as pyruvate) inhibits calcium entry into the cytoplasm in astrocytes and neurons, and shifts metabolism towards respiration [74]. Lactate also inhibits calcium entry into pancreatic a-cells and glucagon secretion[75]. This is similar to the effect of carbon sources on calcium signaling in yeast that we have determined, and it is thus possible that a similar universal carbon source sensing role for calcium is conserved from yeast to man. In humans, calcium signaling is implicated in many disease states, such as diabetes [72,73,75], Alzheimers disease [76] and cancer [77].

To summarize, we have expanded the scope of calcium signaling to be a pan-carbon source sensor, with a different analogue response to each carbon source. We have also shown that a broader range of genes are regulated by calcium signaling than previously determined – both positively (*ZWF1*) and negatively (*ADH2*). The ratio of G6P/G1P is converted into a calcium signal, possibly via regulation of the cytoplasmic pH by Pma1. Furthermore, prior calcium signaling events serves as an **ionic memory** that impinges upon future carbon source events.

## Acknowledgements

Many thanks to all members of the Kupiec Lab for fruitful discussions, and to Roy Salinas for support. Many thanks to the Dequin, Young, Arino, Berger, Cocetti, Cannon and Karpov labs for providing strains and plasmids. This work was supported by grants from the Israel Science Foundation and the Minerva Stiftung to M.K.

## Author Contributions

Conceptualization - KSL and MK; Methodology - KSL; Investigation - KSL; Writing – Original Draft, KSL and MK; Writing – KSL and MK; Funding Acquisition - MK; Resources - MK; Supervision – MK.

## Materials and Methods

### CONTACT FOR REAGENT AND RESOURCE SHARING

Further information and requests for resources and reagents should be directed to and will be fulfilled by the Lead Contact, Martin Kupiec. There are no restrictions on access to strains and plasmids.

### EXPERIMENTAL MODEL AND SUBJECT DETAILS

Experiments were conducted using *S. cerevisiae*. DH5a bacteria were used for plasmid propagation and standard procedures. Strains used are listed in **Table 1**; plasmids used are listed in **Table 2**. All strains were related to the industrial champagne derived yeast haploid V5 [78] except for strains used in **Figure 7** which are in the BY4741a background [79] Yeasts were transformed with DNA using the frozen lithium acetate method [79]. Deletions were confirmed by PCR. Selective markers in plasmids were exchanged by gap repair [80].

**Table 1:**
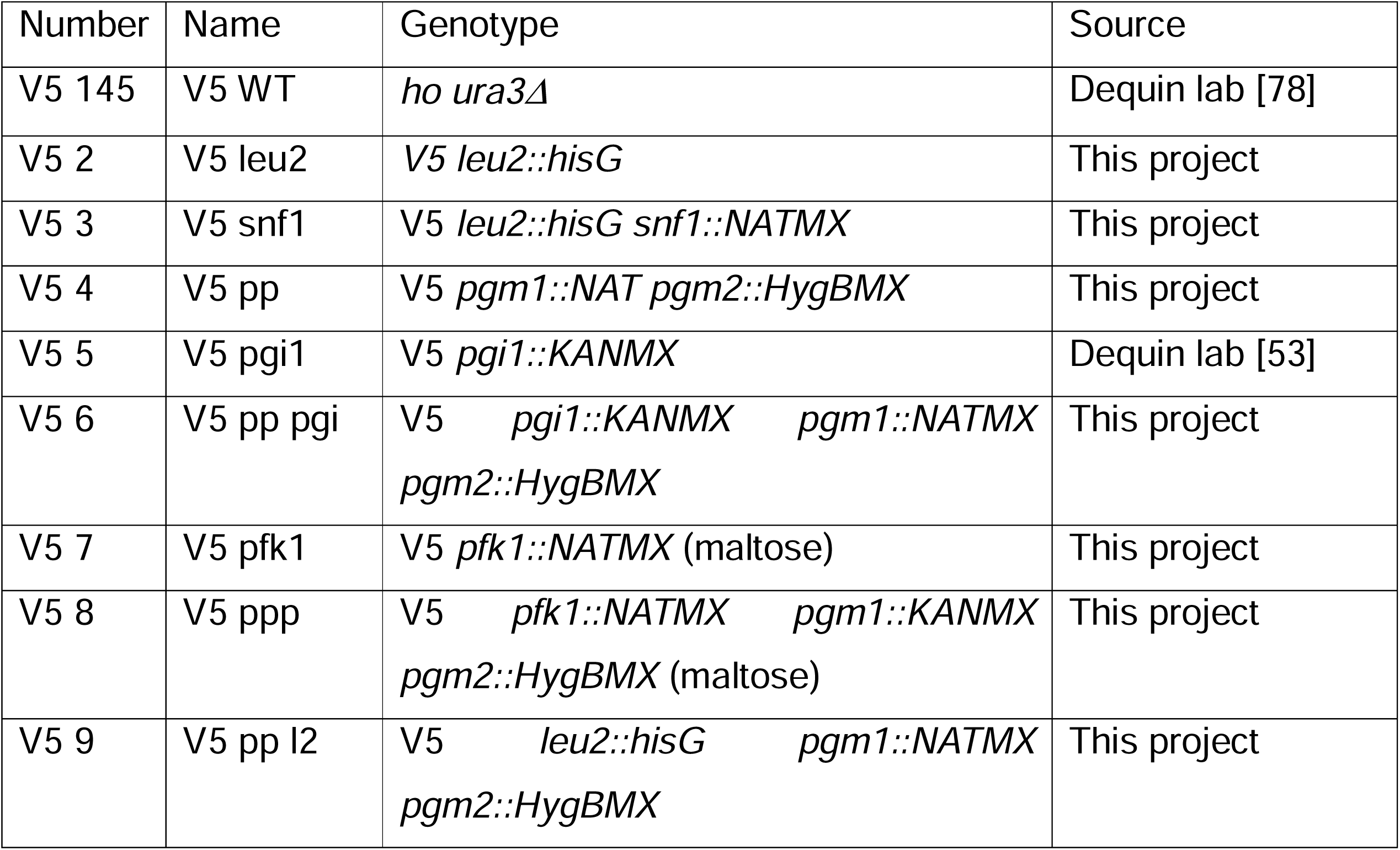
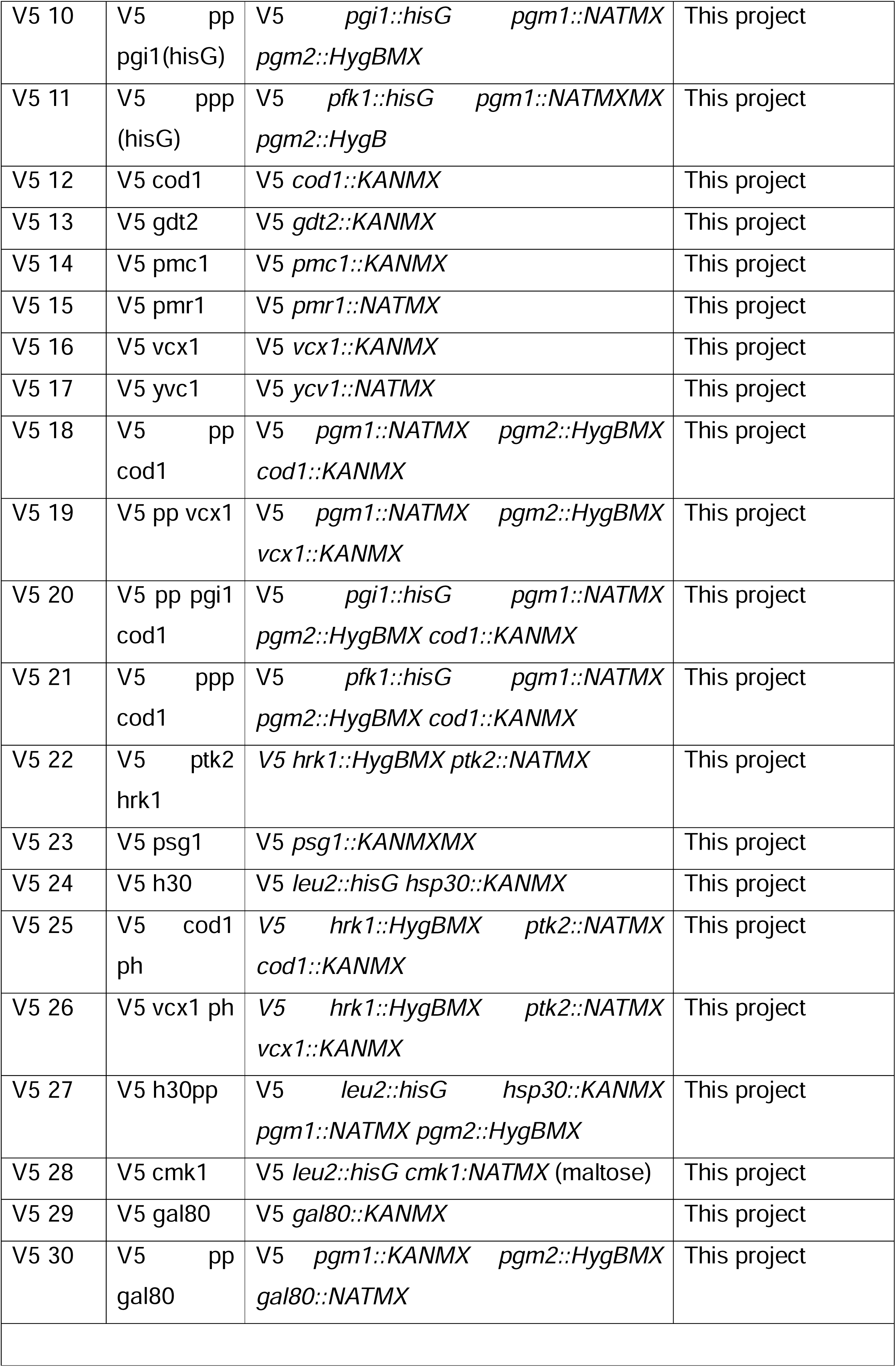
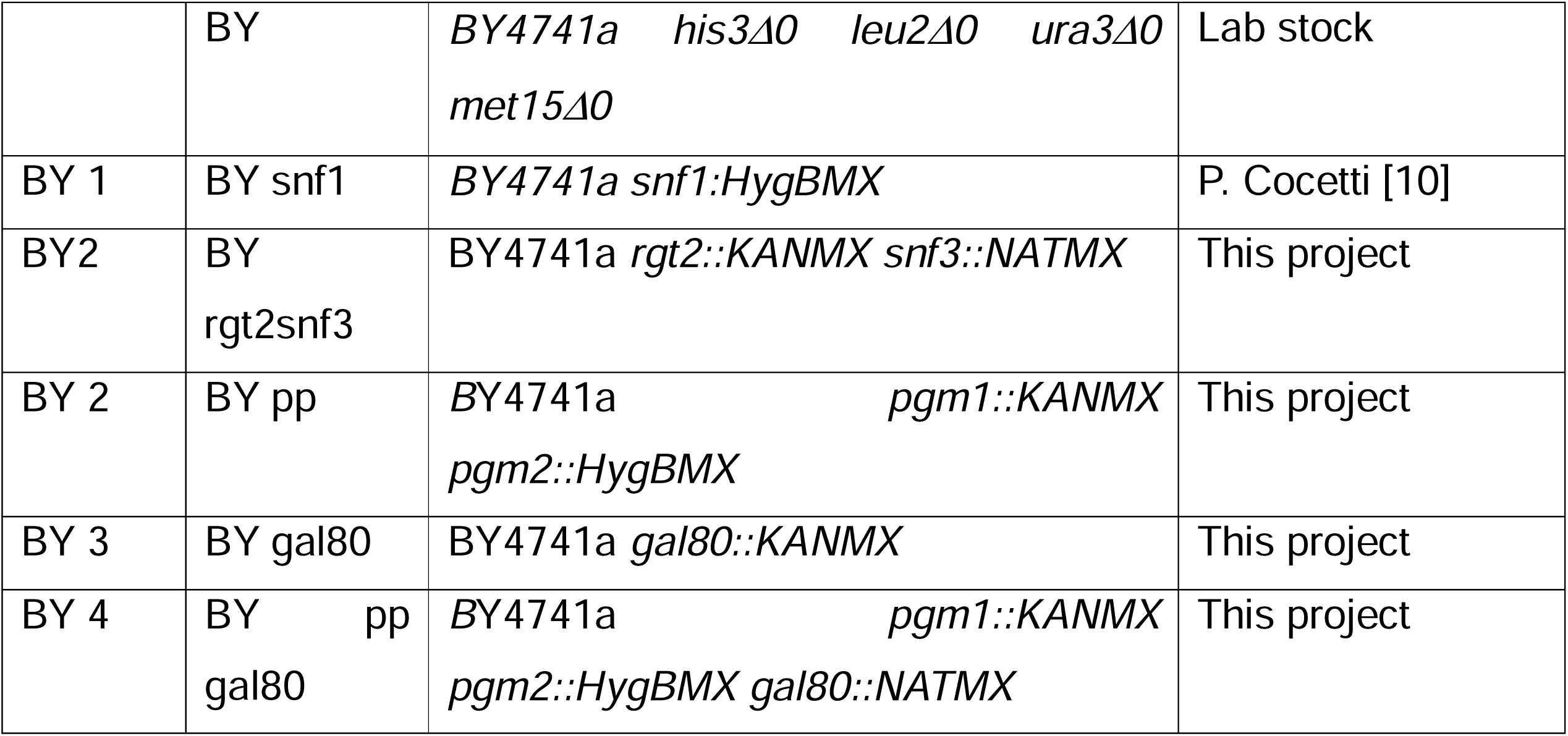
Yeast Strains.

| Number | Name | Genotype | Source |
| --- | --- | --- | --- |
| V5 145 | V5 WT | *ho ura3Δ* | Dequin lab [78] |
| V5 2 | V5 leu2 | *V5 leu2::hisG* | This project |
| V5 3 | V5 snf1 | *V5 leu2::hisG snf1::NATMX* | This project |
| V5 4 | V5 pp | *V5 pgm1::NAT pgm2::HygBMX* | This project |
| V5 5 | V5 pgi1 | *V5 pgi1::KANMX* | Dequin lab [53] |
| V5 6 | V5 pp pgi | *V5 pgi1::KANMX pgm1::NATMX*<br>*pgm2::HygBMX* | This project |
| V5 7 | V5 pfk1 | *V5 pfk1::NATMX* (maltose) | This project |
| V5 8 | V5 ppp | *V5 pfk1::NATMX pgm1::KANMX*<br>*pgm2::HygBMX* (maltose) | This project |
| V5 9 | V5 pp l2 | *V5 leu2::hisG pgm1::NATMX*<br>*pgm2::HygBMX* | This project |
| V5 10 | V5 pp<br>pgi1(hisG) | V5 *pgi1::hisG* *pgm1::NATMX*<br>*pgm2::HygBMX* | This project |
| V5 11 | V5 ppp<br>(hisG) | V5 *pfk1::hisG* *pgm1::NATMXMX*<br>*pgm2::HygB* | This project |
| V5 12 | V5 cod1 | V5 *cod1::KANMX* | This project |
| V5 13 | V5 gdt2 | V5 *gdt2::KANMX* | This project |
| V5 14 | V5 pmc1 | V5 *pmc1::KANMX* | This project |
| V5 15 | V5 pmr1 | V5 *pmr1::NATMX* | This project |
| V5 16 | V5 vcx1 | V5 *vcx1::KANMX* | This project |
| V5 17 | V5 yvc1 | V5 *ycv1::NATMX* | This project |
| V5 18 | V5 pp<br>cod1 | V5 *pgm1::NATMX* *pgm2::HygBMX*<br>*cod1::KANMX* | This project |
| V5 19 | V5 pp vcx1 | V5 *pgm1::NATMX* *pgm2::HygBMX*<br>*vcx1::KANMX* | This project |
| V5 20 | V5 pp pgi1<br>cod1 | V5 *pgi1::hisG* *pgm1::NATMX*<br>*pgm2::HygBMX* *cod1::KANMX* | This project |
| V5 21 | V5 ppp<br>cod1 | V5 *pfk1::hisG* *pgm1::NATMX*<br>*pgm2::HygBMX* *cod1::KANMX* | This project |
| V5 22 | V5 ptk2<br>hrk1 | V5 *hrk1::HygBMX* *ptk2::NATMX* | This project |
| V5 23 | V5 psg1 | V5 *psg1::KANMXMX* | This project |
| V5 24 | V5 h30 | V5 *leu2::hisG* *hsp30::KANMX* | This project |
| V5 25 | V5 cod1<br>ph | V5 *hrk1::HygBMX* *ptk2::NATMX*<br>*cod1::KANMX* | This project |
| V5 26 | V5 vcx1 ph | V5 *hrk1::HygBMX* *ptk2::NATMX*<br>*vcx1::KANMX* | This project |
| V5 27 | V5 h30pp | V5 *leu2::hisG* *hsp30::KANMX*<br>*pgm1::NATMX* *pgm2::HygBMX* | This project |
| V5 28 | V5 cmk1 | V5 *leu2::hisG* *cmk1::NATMX* (maltose) | This project |
| V5 29 | V5 gal80 | V5 *gal80::KANMX* | This project |
| V5 30 | V5 pp<br>gal80 | V5 *pgm1::KANMX* *pgm2::HygBMX*<br>*gal80::NATMX* | This project |
|  | BY | *BY4741a his3Δ0 leu2Δ0 ura3Δ0 met15Δ0* | Lab stock |
| BY 1 | BY snf1 | *BY4741a snf1::HygBMX* | P. Cocetti [10] |
| BY2 | BY<br>rgt2snf3 | *BY4741a rgt2::KANMX snf3::NATMX* | This project |
| BY 2 | BY pp | *BY4741a pgm1::KANMX*<br>*pgm2::HygBMX* | This project |
| BY 3 | BY gal80 | *BY4741a gal80::KANMX* | This project |
| BY 4 | BY pp<br>gal80 | *BY4741a pgm1::KANMX*<br>*pgm2::HygBMX gal80::NATMX* | This project |

**Table 2.** Plasmids.

| Number | Backbone | Contains | Original Backbone | Source |
| --- | --- | --- | --- | --- |
| 85 | pRS 315 | *prSNF1::SNF1-2HA* |  | Berger lab [81] |
| 319 | pRS 316 | *4x prCDRE::LacZ* | pAMS366 | Arino lab [55] |
| 411 | pRS 316 | *prZWF1::LacZ* |  | Karpov lab [42] |
| 467 | pRS 315 | *prRAS2::Ras2<sup>G19V</sup>* | pRS 316 | Cannon lab [45] |
| 488 | pRS 316 | *prADH1:ADR1<sup>S230A</sup>* | pRS 314 | Young lab [43] |
| 919 | pRS 316 | *prADH2::LacZ* |  | Young lab [43] |
| 1367 | pRS 315 | *prPMA1 :: Pma1Δ901* |  | Kupiec lab [62] |

#### β-galactosidase assay experimental details for Figures 1-6

##### *ADH2* expression

Yeast containing a plasmid expressing β-galactosidase from an *ADH2* promoter were grown overnight at 30°C in selective media containing 4% glucose, diluted and then grown for another 3 hours. When experiments involved cells deleted for *PGI1*, all strains were grown in media containing 2% Fructose with 0.05% Glucose [53]. Samples were taken for determining *ADH2* expression under repressive conditions (t=0). Cells were washed three times with ddH_2_0 and then resuspended in selective media lacking any carbon source. 10mM MgCl_2_ and FK506 (from a 20mg/ml stock in 90% Ethanol/10% Tween-20) was added where indicated. Cells were aliquoted into tubes, and the different carbon sources added to give a final concentration of 2% for glucose, fructose, sucrose, maltose, galactose and ethanol; and 3% for glycerol. Cells were grown for a further three hours and then *ADH2* expression determined by β-galactosidase assay (t=3). Experiments were conducted in triplicate.

##### *ZWF* expression

V5 Yeast containing a plasmid expressing β-galactosidase from a *ZWF1* promoter (358 bp upstream of the ATG) were grown overnight at 30°C in selective media containing 4% glucose, diluted and then grown for another 3 hours. When experiments involved cells deleted for *PGI1*, all strains were grown in media containing 2% Fructose with 0.05% Glucose [53]. Samples were taken for determining *ZWF1* expression under basal conditions (t=0). Cells were washed three times with ddH_2_0 and then resuspended in selective media lacking any carbon source. MMS was added to give a final concentration of 0.05%. 10mM MgCl_2_ and FK506 (from a 20mg/ml stock in 90% Ethanol/10% Tween-20) was added where indicated. Cells were aliquoted into tubes, and the different carbon sources added to give a final concentration of 2% for glucose, fructose, sucrose, maltose, galactose and ethanol; and 3% for glycerol. Cells were grown for a further three hours and then *ZWF1* expression determined by β-galactosidase assay (t=3). Experiments were conducted in triplicate. Note that for experiment of **Figure S1D** BY4741a yeast were used and cells grown overnight and the next morning in 2% ethanol, in order to prevent suppressor mutations occurring in the *rgt2Δ snf3Δ* cells.

##### *CDRE* expression

**Figure 2E**. Wild-type V5 yeast containing a plasmid expressing β-galactosidase from 26 bp fragment of the *FKS2* promoter (CDRE motif) [55] were grown overnight at 30°C in selective media containing 2% of the carbon source indicated (glycerol at 3%), with or without the calcium chelator EGTA (1mM). Cells were diluted in the morning into the same carbon sources (and more EGTA added where required) and grown for an additional 3 hours before the basal *CDRE* expression was determined by β-galactosidase assay. Experiments were conducted in triplicate. **Figures 2F, S2A, S2B, S2C, 4A, S4A, 5A, 5D, S5A, 6A**. Indicated V5 cells containing a plasmid expressing β-galactosidase from 26bp fragment of the *FKS2* promoter (CDRE motif) [55] were grown overnight at 30°C in selective media containing 3% glycerol + 0.1% glucose. 1mM EGTA and FK506 (from a 20mg/ml stock in 90% Ethanol/10% Tween-20) was present where indicated. Cells were aliquoted into tubes and diluted in the morning into media containing 3% glycerol (no glucose) and grown for 3 hours. Samples were taken for determining *CDRE* expression under basal conditions (t=0). Indicated carbon sources (UPPERCASE) were added to 2% and the cells grown for 90 minutes before *CDRE* expression was determined by β-galactosidase assay (t=1.5). Experiments were conducted in triplicate. Note that this method cannot be used for *pgi1Δ* cells, so only experiments using *pfk1Δ* cells were performed to block glycolysis.

**Figure 7 and S6** experiments.

Calcium accumulation in *pgm1Δpgm2Δ* [39,48] or 15mM LiCl treated [40,65] cells is due to Galactose-1-Phosphate accumulation and this prevents growth in BY4741 cells [48], (**Figure S6A**). However, V5 cells are far less capable than BY4741 cells to utilize galactose as a carbon source (**Figure S6A**) but unlike in the BY4741 background V5 *pgm1Δ pgm2Δ* cells are viable on galactose. (This deficiency in galactose metabolism was not rescued by deletion of *GAL80* (**Figure S6A**) encoding a repressor of GAL gene expression (61) . We therefore conducted this set of experiments in the BY4741 background.

The aim of these experiments was to use either inhibition of deletion of phosphoglucomutase (PGM) activity to cause the cells to accumulate calcium when grown in galactose [40,65] which is then released by addition of glucose For Figure 7A (V5 cells) and **Figure S6B** (BY4741 cells), cells were grown overnight in the indicated carbon source and diluted in the morning with the same carbon source and allowed to grow for an additional three hours. The basal *CDRE* expression was determined by β-galactosidase assay. Then 2% glucose was added and after 90 mins the *CDRE* expression was determined by β-galactosidase assay (Figure 7B for V5 cells, **Figure S6C** for BY4741 cells). ). Experiments were conducted in triplicate.

##### Figures 7C, D

BY4741 cells expressing β-galactosidase from either a *CDRE* (Figure 7C) or a *ZWF1* (Figure 7D) promoter were grown overnight at 30°C in selective media containing 3% glycerol + 0.1% glucose. Cells were diluted 5-fold and grown for 6 hours in media containing 3% glycerol (no glucose). WT cells to be treated with 15mM LiCl and to be grown in media containing 2% galactose or *pgm1Δpgm2Δ*cells to be grown in media containing 2% galactose were diluted 4x. Other cells were diluted 50x. Cells were distributed into tubes and grown under indicated conditions (combinations of 2% glucose, 2% galactose, 15mM LiCl, 1mM EGTA (Figure 7C)) for 16 hours and the basal β-galactosidase activity measured (t=0). For Figure 7D (*ZWF1* expression), 0.05% MMS was now added. The glucose grown cells now had another 2% glucose added to them (glucose-GLUCOSE), the galactose cells now had either 2% glucose (galactose-GLUCOSE) or 2% galactose (galactose-GALACTOSE) added. 10mM MgCl2 was added as indicated (Figure 7D). After 90 minutes (Figure 7C) or 3 hours (Figure 7D) the β-galactosidase activity was determined. Note that all the WT cells used in each experiment were from the same colony grown in 3% glycerol + 0.1% glucose Experiments were performed in triplicate.

##### β-galactosidase assay procedure

Cell concentration was determined by reading 100μl of cells at 595nm. 20μl of cells was added to the β-galactosidase reaction mix for *ADH2* and *CDRE* and 50μl for *ZWF1* expression determination (40μl YPER (Pierce 78990), 80μl Z-buffer (120mM Na_2_HPO_4_, 80mM NaH_2_PO_4_, 20mM KCl, 2mM MgSO_4_), 24μl ONPG (4mg/ml), 0.4μl β-mercaptoethanol) and incubated at 30°C for 15 minutes for *ADH2* and *CDRE* expression and for 30 minutes for *ZWF1* expression determination. Reactions were stopped by addition of 50μl 1.08M Na_2_CO_3_. The Eppendorf tubes were centrifuged for 1 minute at full speed to pellet the cell debris, and 200μl supernatant was removed, and absorbance read at 415nm using a microplate reader. Miller Units were calculated by the equation Miller Units = (1000*A_415_)/(time*volume of cells*A_595_-0.055, where the A_415_ and A_595_ have been corrected for blanking and path length (final path length = 1cm). Three biological replicates were measured. Error bars are +/- 1 standard deviation.

### Western Blot experimental details

Indicated V5 *snf1Δ* cells Figure 1B) with a plasmid expressing Snf1-2HA from a Snf1 promoter in pRS315 (or a wild-type strain with pRS315) were grown in selective media containing 4% glucose overnight. Cells were washed twice with 25ml ddH_2_0 and resuspended in selective media lacking any carbon source. Cells were aliquoted into glass tubes and indicated carbon source added to 2% (glycerol to 3%). Cells were grown at 30^◦^C for 15 minutes before processing.

For **Figure S1C**, WT V5 cells were grown in selective media containing 4% glucose overnight. Cells were washed twice with 25ml ddH_2_0 and resuspended in selective media lacking any carbon source. Cells were aliquoted into glass tubes and indicated carbon source added to 2% (glycerol to 3%). Cells were grown at 30^◦^C for 3 hours before processing.

Cells were harvested and protein solubilized in sample buffer by the method developed by the Kuchin group to prevent activation of Snf1 by centrifugation [82]. Cells were boiled for 5 minutes before treatment with 0.2M NaOH. Sample buffer volume was adjusted to give equal OD for all samples. After running samples on 10% polyacrylamide gels, the proteins were wet transferred to nitrocellulose membranes. Antibodies used were mouse α-Pgk1 (Abcam 113687) for a loading control, mouse anti-HA (Abcam), mouse anti-polyhistidine (for detecting native Snf1 via the 13 polyhisitidine tract in the PKR of Snf1 [62]) (Abcam) and rabbit α-phospo-T172 (AMPK) (Cell Signalling Technology), all at 1/1000 dilution. Since Snf1 runs at 72kDa the membrane was cut between 55 and 65 kDa, the lower half used for Pgk1 and the upper for HA, polyHIStidine or phospho-T172. Secondary antibodies were conjugated to HRP. Images were minimally processed using ImageJ.

### Serial dilution experiment details

Freshly grown cells were taken from plates and resuspended in ddH_2_0. Following determination of OD_600_ the concentration of the yeast was normalized to OD4/ml. 30μl was taken and serially diluted ten-fold 5 times, and 5μl final four dilutions spotted onto indicated plates. Photographs were taken after 2 days (**S1B**, **S6A**) as indicated.

## QUANTIFICATION AND STATISTICAL ANALYSIS

Statistical details are to be found in the figure legends. For β-galactosidase and Western Blots, N represents the number of biological repeats. Error bars are one standard deviation. Western Blots were quantified by finding the ratio of Snf1:Pgk1 (or phospho-Snf1:Pgk1) and then the ratio of [phospho-Snf1:Pgk1]:[Snf:Pgk1] for each experiment. Since the intensity of Pgk1 was not consistent over multiple experiments, these measurements were then normalized with WT ethanol representing 100% phosphorylation.

## KEY RESOURCES TABLE

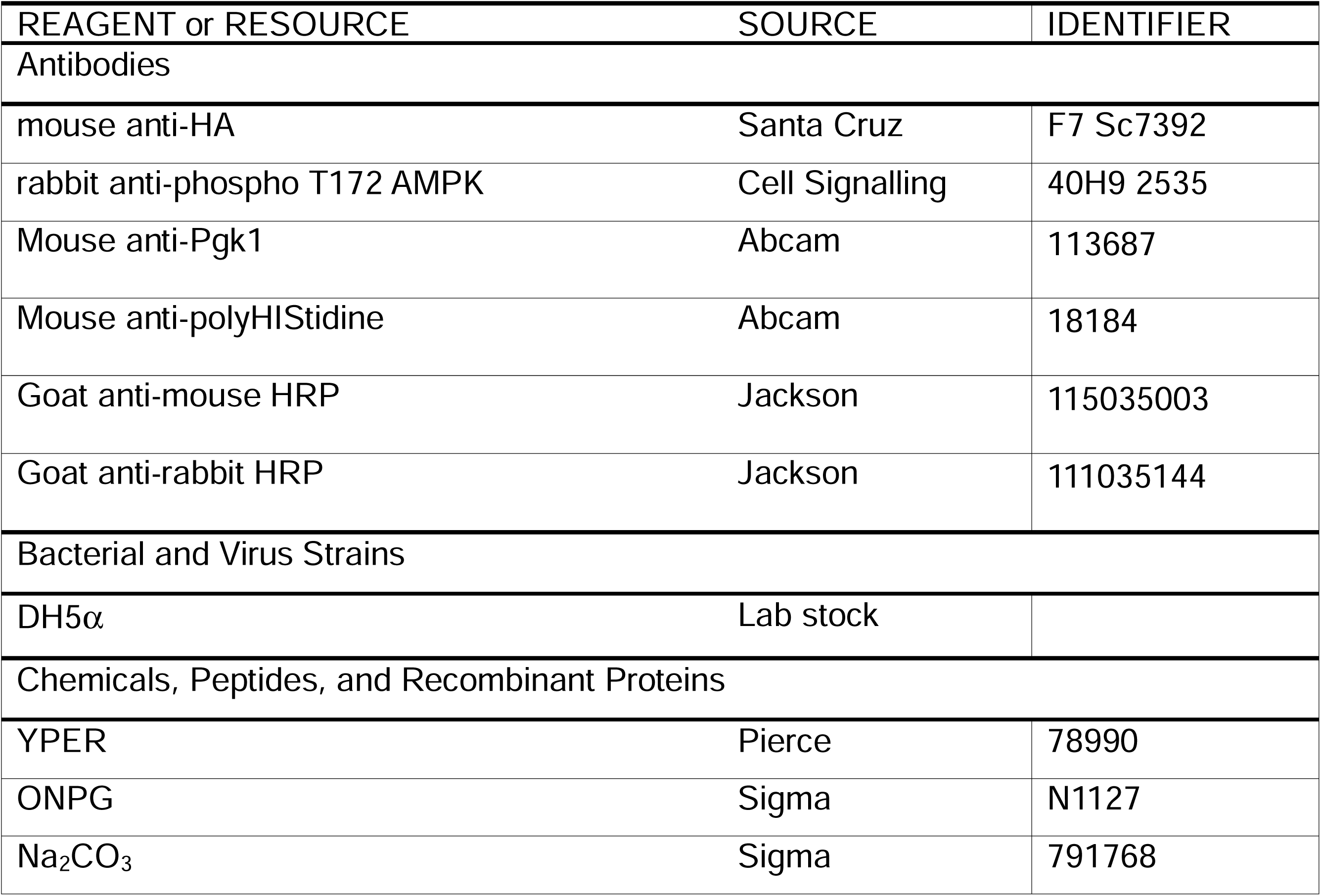
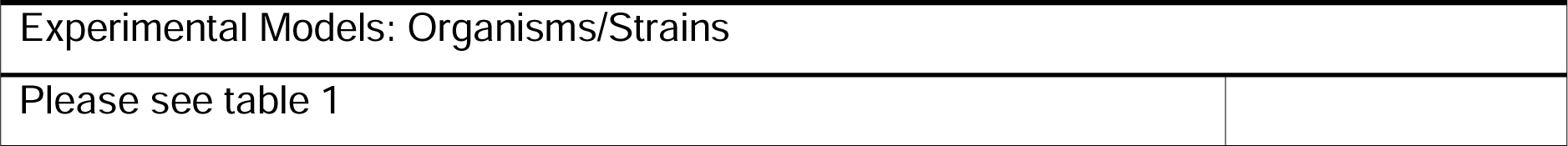

## Supplemental Figure Legends

**Figure S1 (pertaining to Figure 1).**
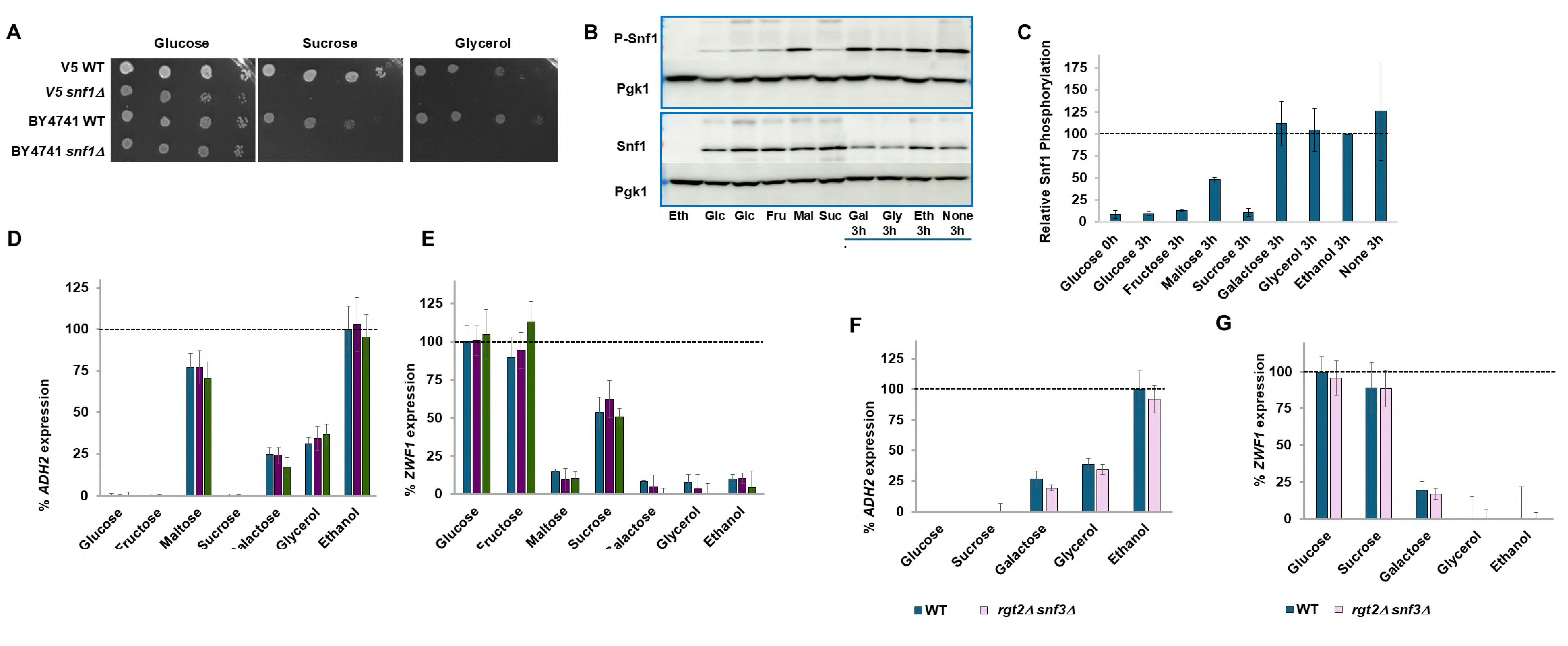
**B**. Serial dilution assay performed as described in materials and methods to show that *snf1Δ* cells in both BY4741 and V5 backgrounds do not grow on sucrose nor on glycerol as their carbon source. **C**. Glucose-grown cells were washed twice with ddw and then grown for 3 hours in the indicated carbon source (**Glu**cose, **Fru**ctose, **Mal**tose, **Suc**rose, **Gal**actose, **Gly**cerol, **Eth**anol or **None**) as described in Materials and Methods. Cells were processed and Western Blots obtained as described in Materials and Methods. The experiment was repeated **three** times. Pgk1 is used as a loading control. P-Snf1 is Snf1 phosphorylated at T210. Anti-PolyHIStidine was used to detect native Snf1. **D**. Quantifications of Western Blots of **Figure 1A** were performed using ImageJ. **E.** V5 cells were grown overnight in 4% glucose, diluted and grown the following morning for an additional three hours. Basal *ADH2* expression was determined by β-galactosidase assay. Cells were washed and resuspended in media lacking any carbon source, and the indicated carbon sources added. *ADH2* expression was determined after three hours of growth at 30^◦^C. Expression rate is normalized to WT in ethanol. N=3**. F.** V5 cells were grown overnight in 4% glucose, diluted and grown the following morning for an additional three hours. Basal *ZWF1* expression was determined by β-galactosidase assay. Cells were washed and resuspended in media lacking any carbon source, and the indicated carbon sources with 0.05% MMS. *ZWF1* expression was determined after three hours of growth at 30^◦^C. Expression rate is normalized to WT in glucose. N=3**. G**. BY4741 cells were grown overnight in 4% glucose, diluted and grown the following morning for an additional three hours. Basal *ADH2* expression was determined by β-galactosidase assay. Cells were washed and resuspended in media lacking any carbon source, and the indicated carbon sources added. *ADH2* expression was determined after three hours of growth at 30^◦^C. Expression rate is normalized to WT in ethanol. N=3**. H.** BY4741 cells were grown overnight in 2% ethanol, diluted and grown the following morning for an additional three hours. Basal *ZWF1* expression was determined by β-galactosidase assay. Cells were washed and resuspended in media lacking any carbon source, and the indicated carbon sources with 0.05% MMS. *ZWF1* expression was determined after three hours of growth at 30^◦^C. Expression rate is normalized to WT in glucose. N=3.

**Figure S2 (pertaining to Figure 2).**
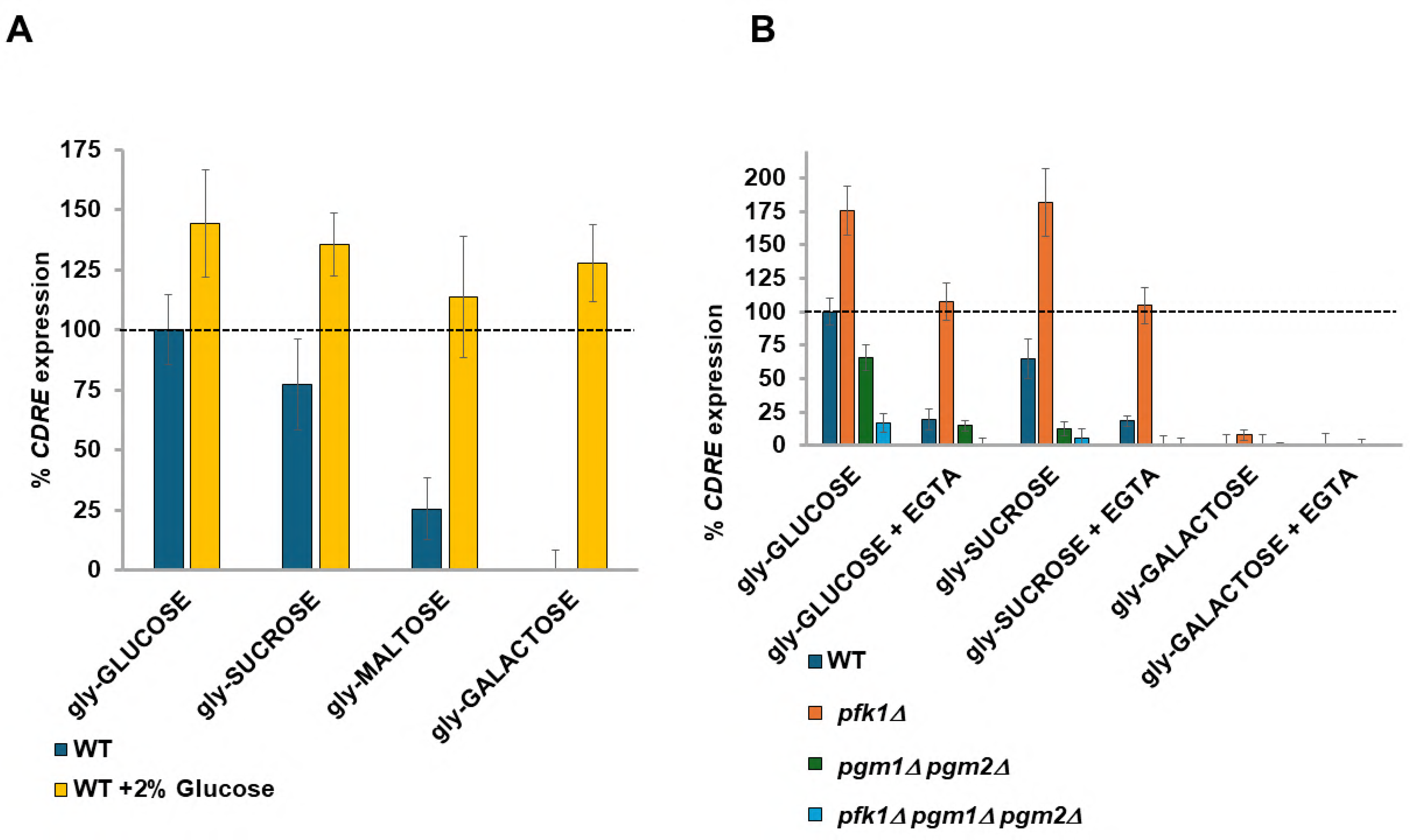
**A.** V5 cells were grown overnight at 30°C in selective media containing 3% glycerol + 0.1% glucose. Cells were aliquoted into tubes and diluted in the morning into media containing 3% glycerol (no glucose) and grown for 3 hours. Samples were taken for determining *CDRE* expression under basal conditions (t=0). Indicated carbon sources (UPPERCASE) (also with additional 2% glucose where indicated) were added to 2% and the cells grown for 90 minutes before *CDRE* expression was determined by β-galactosidase assay (t=1.5). Expression is normalized to WT in glucose. N=3. **B**. V5 cells were grown overnight at 30°C in selective media containing 3% glycerol + 0.1% glucose. Cells were aliquoted into tubes and diluted in the morning into media containing 3% glycerol (no glucose) and grown for 3 hours. Samples were taken for determining *CDRE* expression under basal conditions (t=0). Indicated carbon sources (UPPERCASE) were added to 2% and 1mM EGTA where indicated and the cells grown for 90 minutes before *CDRE* expression was determined by β-galactosidase assay (t=1.5). Expression is normalized to WT in glucose. N=3.

**Figure S3 (pertaining to Figure 3).**
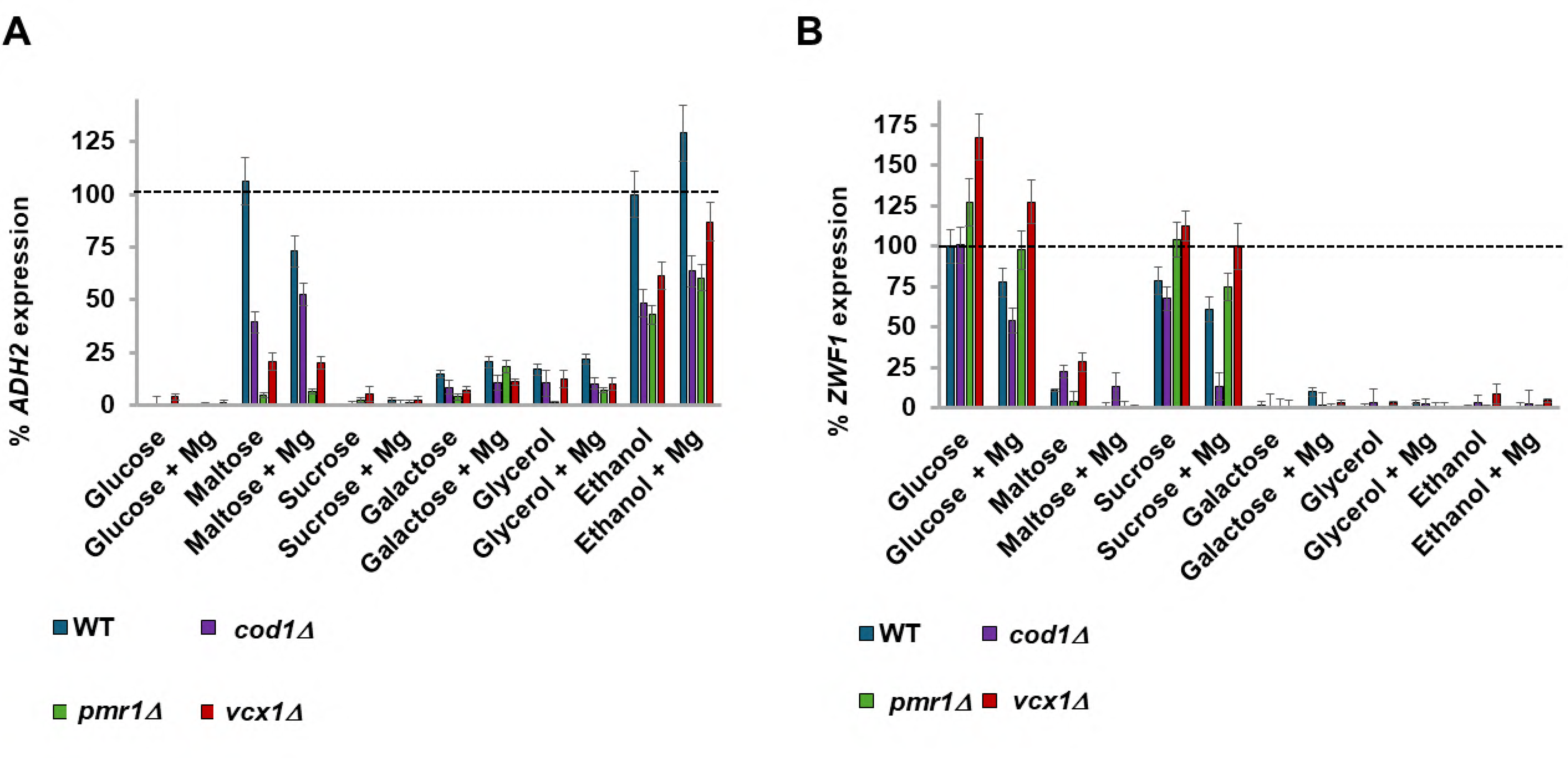
**A.** V5 cells were grown overnight in 4% glucose, diluted and grown the following morning for an additional three hours. Basal *ADH2* expression was determined by β-galactosidase assay. Cells were washed and resuspended in media lacking any carbon source, and the indicated carbon sources added with 10mM MgCl_2_ where indicated. *ADH2* expression was determined after three hours of growth at 30^◦^C. Expression rate is normalized to WT in ethanol. N=3. B. V5 cells were grown overnight in 4% glucose, diluted and grown the following morning for an additional three hours. Basal *ZWF1* expression was determined by β-galactosidase assay. Cells were washed and resuspended in media lacking any carbon source, and the indicated carbon sources with 0.05% MMS. 10mM MgCl_2_ was added where indicated. *ZWF1* expression was determined after three hours of growth at 30^◦^C. Expression rate is normalized to WT in glucose. N=3.

**Figure S4 (pertaining to Figure 4).**
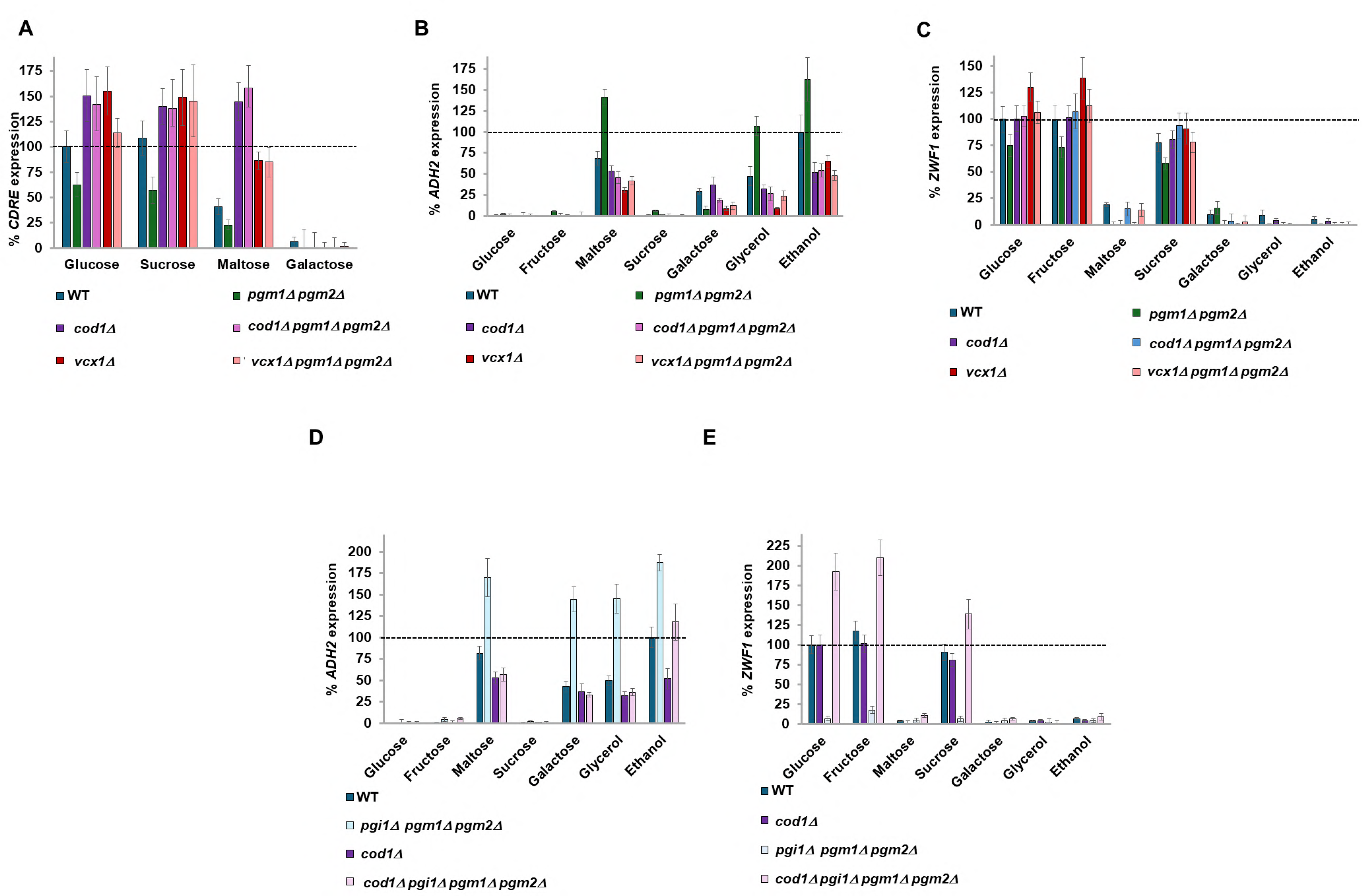
**A.** V5 cells were grown overnight at 30°C in selective media containing 3% glycerol + 0.1% glucose. Cells were aliquoted into tubes and diluted in the morning into media containing 3% glycerol (no glucose) and grown for 3 hours. Samples were taken to determine *CDRE* expression under basal conditions (t=0). Indicated carbon sources (UPPERCASE) were added to 2% and the cells grown for 90 minutes before *CDRE* expression was determined by β-galactosidase assay (t=1.5). Expression is normalized to WT in glucose. N=3. **B, D**. V5 cells were grown overnight in 4% glucose (**B**) or 2% Fructose + 0.05% glucose (**D**), diluted and grown the following morning for an additional three hours. Basal *ADH2* expression was determined by β-galactosidase assay. Cells were washed and resuspended in media lacking any carbon source, and the indicated carbon sources added. *ADH2* expression was determined after three hours of growth at 30^◦^C. Expression rate is normalized to WT in ethanol. N=3**. C, E.** V5 cells were grown overnight in 4% glucose (**C**) or 2% Fructose + 0.05% glucose (**E**), diluted and grown the following morning for an additional three hours. Basal *ZWF1* expression was determined by β-galactosidase assay. Cells were washed and resuspended in media lacking any carbon source, and the indicated carbon sources with 0.05% MMS. *ZWF1* expression was determined after three hours of growth at 30^◦^C. Expression rate is normalized to WT in glucose. N=3.

**Figure S5 (pertaining to Figure 5).**
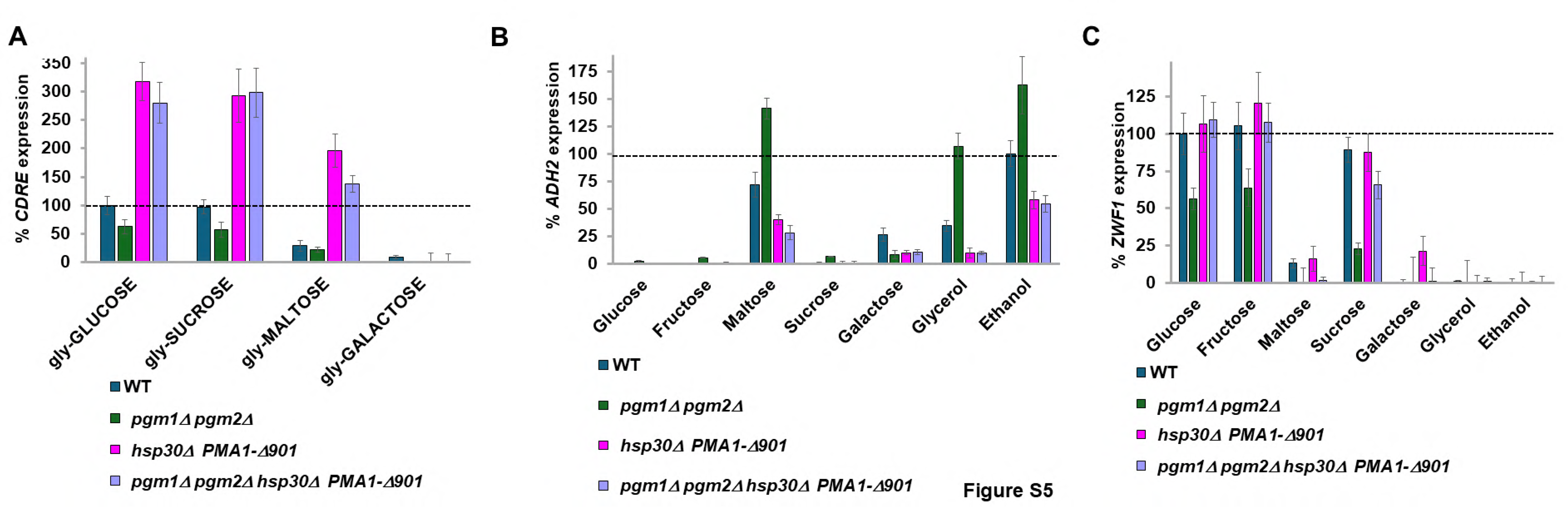
**A**. V5 cells were grown overnight at 30°C in selective media containing 3% glycerol + 0.1% glucose. Cells were aliquoted into tubes and diluted in the morning into media containing 3% glycerol (no glucose) and grown for 3 hours. Samples were taken to determine *CDRE* expression under basal conditions (t=0). Indicated carbon sources (UPPERCASE) were added to 2% and the cells grown for 90 minutes before *CDRE* expression was determined by β-galactosidase assay (t=1.5). Expression is normalized to WT in glucose. N=3. **B**. V5 cells were grown overnight in 4% glucose, diluted and grown the following morning for an additional three hours. Basal *ADH2* expression was determined by β-galactosidase assay. Cells were washed and resuspended in media lacking any carbon source, and the indicated carbon sources added. *ADH2* expression was determined after three hours of growth at 30^◦^C. Expression rate is normalized to WT in ethanol. N=3**. C.** V5 cells were grown overnight in 4% glucose, diluted and grown the following morning for an additional three hours. Basal *ZWF1* expression was determined by β-galactosidase assay. Cells were washed and resuspended in media lacking any carbon source, and the indicated carbon sources with 0.05% MMS. *ZWF1* expression was determined after three hours of growth at 30^◦^C. Expression rate is normalized to WT in glucose. N=3.

**Figure S6 (pertaining to Figure 7).**
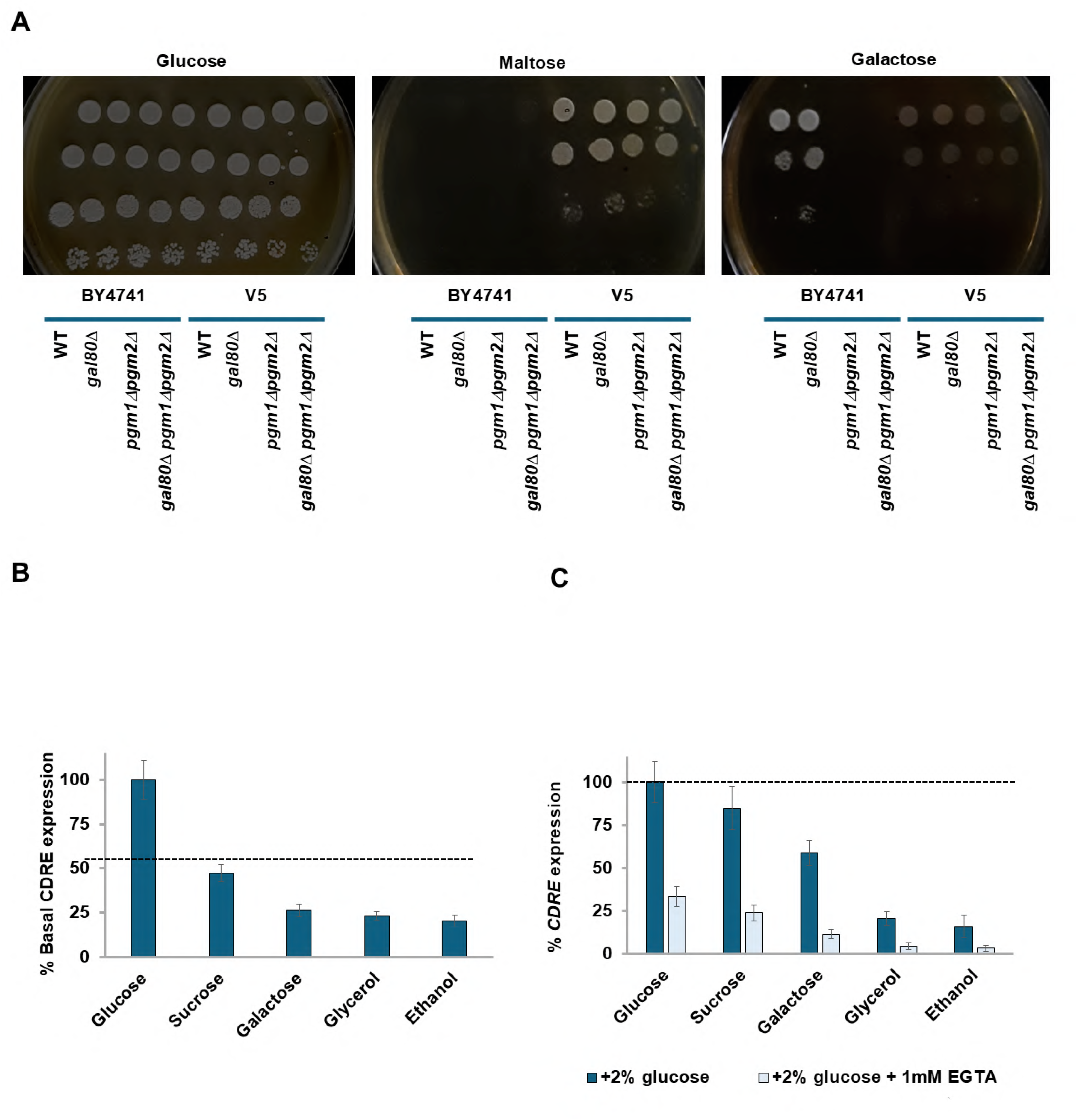
**A.** Serial dilution assay showing growth of indicated cells on indicated media. Cells were treated as described in Materials and Methods. **B.** Wild-type BY4741 yeast were grown overnight at 30°C in selective media containing 2% of the carbon source indicated (glycerol at 3%), with or without the calcium chelator EGTA (1mM). Cells were diluted in the morning into the same carbon sources (and more EGTA added where required) and grown for an additional 3 hours before the basal *CDRE* expression was determined by β-galactosidase assay. Expression is normalized to WT in glucose. N=3. **C**. 2% glucose or 2% glucose + 1mM EGTA was added to the cells from **Figure S6A**. and the cells grown for 90 minutes before *CDRE* expression was determined by β-galactosidase assay (t=1.5). Expression is normalized to WT in glucose. N=3.

---

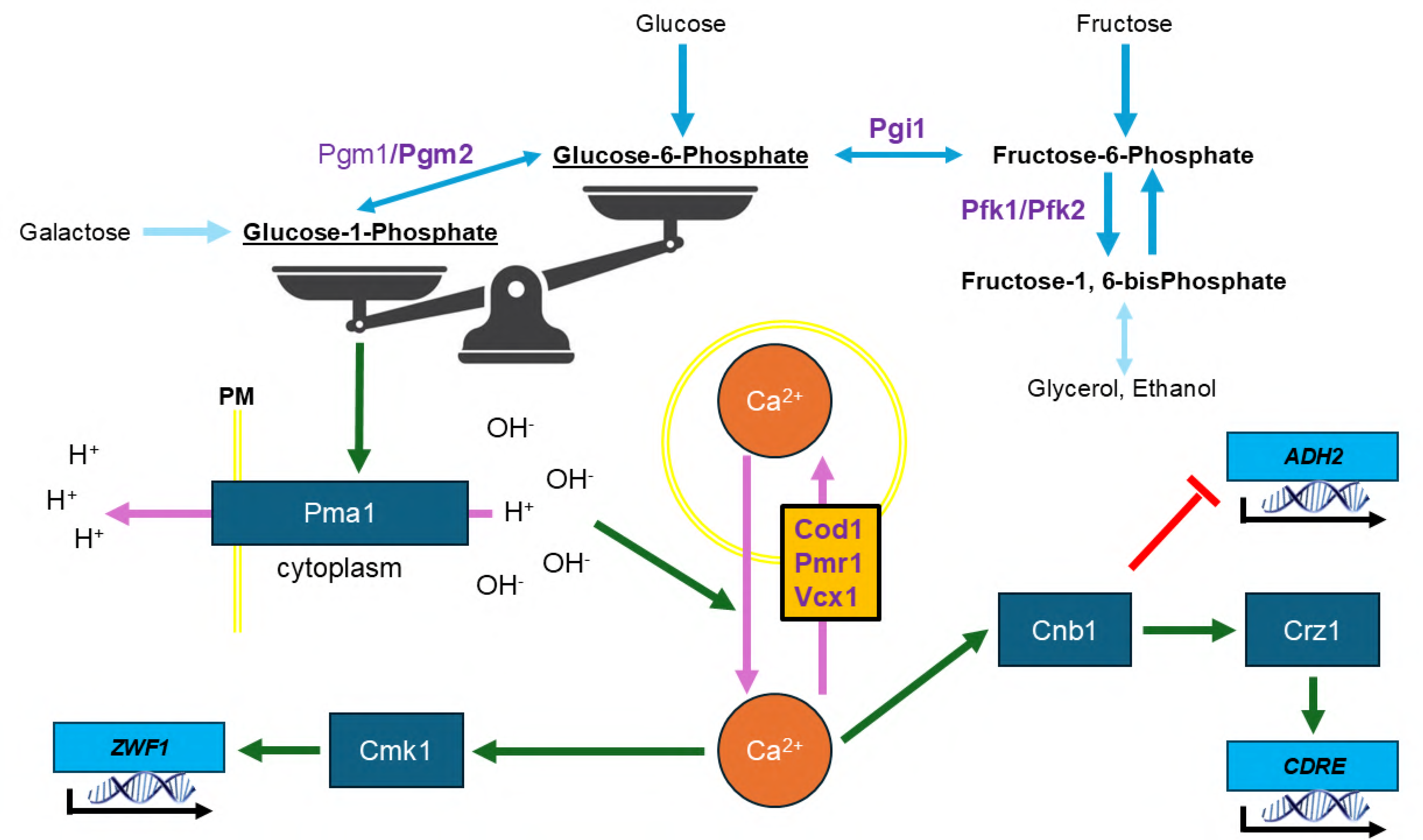

## References

1. Simpson-Lavy, K.; Kupiec, M. Carbon catabolite repression: not only for glucose. Curr Genet 2019, 65, 1321–1323, doi:10.1007/s00294-019-00996-6.

2. Gadura, N.; Robinson, L.C.; Michels, C.A. Glc7-Reg1 phosphatase signals to Yck1,2 casein kinase 1 to regulate transport activity and glucose-induced inactivation of Saccharomyces maltose permease. Genetics 2006, 172, 1427–1439, doi:10.1534/genetics.105.051698.

3. Simpson-Lavy, K.J.; Johnston, M. SUMOylation regulates the SNF1 protein kinase. Proc. Natl. Acad. Sci. U. S. A. 2013, 110, 17432–17437, doi:10.1073/pnas.1304839110.

4. Moriya, H.; Johnston, M. Glucose sensing and signaling in Saccharomyces cerevisiae through the Rgt2 glucose sensor and casein kinase I. Proc. Natl. Acad. Sci. U. S. A. 2004, 101, 1572–1577, doi:10.1073/pnas.0305901101.

5. Kim, J.-H.; Brachet, V.; Moriya, H.; Johnston, M. Integration of transcriptional and posttranslational regulation in a glucose signal transduction pathway in Saccharomyces cerevisiae. Eukaryot. Cell 2006, 5, 167–173, doi:10.1128/EC.5.1.167-173.2006.

6. Kim, J.-H.; Johnston, M. Two glucose-sensing pathways converge on Rgt1 to regulate expression of glucose transporter genes in Saccharomyces cerevisiae. J. Biol. Chem. 2006, 281, 26144– 26149, doi:10.1074/jbc.M603636200.

7. Goossens, A.; La Fuente, N. de; Forment, J.; Serrano, R.; Portillo, F. Regulation of yeast H(+)-ATPase by protein kinases belonging to a family dedicated to activation of plasma membrane transporters. Mol. Cell. Biol. 2000, 20, 7654–7661, doi:10.1128/MCB.20.20.7654-7661.2000.

8. Gancedo, J.M. The early steps of glucose signalling in yeast. FEMS Microbiol Rev 2008, 32, 673– 704, doi:10.1111/j.1574-6976.2008.00117.x.

9. Tamaki, H. Glucose-stimulated cAMP-protein kinase A pathway in yeast Saccharomyces cerevisiae. J. Biosci. Bioeng. 2007, 104, 245–250, doi:10.1263/jbb.104.245.

10. Coccetti, P.; Nicastro, R.; Tripodi, F. Conventional and emerging roles of the energy sensor Snf1/AMPK in Saccharomyces cerevisiae. Microb. Cell 2018, 5, 482–494, doi:10.15698/mic2018.11.655.

11. Rubenstein, E.M.; McCartney, R.R.; Zhang, C.; Shokat, K.M.; Shirra, M.K.; Arndt, K.M.; Schmidt, M.C. Access denied: Snf1 activation loop phosphorylation is controlled by availability of the phosphorylated threonine 210 to the PP1 phosphatase. J. Biol. Chem. 2008, 283, 222–230, doi:10.1074/jbc.M707957200.

12. Shashkova, S.; Wollman, A.J.M.; Leake, M.C.; Hohmann, S. The yeast Mig1 transcriptional repressor is dephosphorylated by glucose-dependent and -independent mechanisms. FEMS Microbiol. Lett. 2017, 364, doi:10.1093/femsle/fnx133.

13. Snowdon, C.; Schierholtz, R.; Poliszczuk, P.; Hughes, S.; van der Merwe, G. ETP1/YHL010c is a novel gene needed for the adaptation of Saccharomyces cerevisiae to ethanol. FEMS Yeast Res. 2009, 9, 372–380, doi:10.1111/j.1567-1364.2009.00497.x.

14. Gasmi, N.; Jacques, P.-E.; Klimova, N.; Guo, X.; Ricciardi, A.; Robert, F.; Turcotte, B. The switch from fermentation to respiration in Saccharomyces cerevisiae is regulated by the Ert1 transcriptional activator/repressor. Genetics 2014, 198, 547–560, doi:10.1534/genetics.114.168609.

15. Kim, M.S.; Cho, K.H.; Park, K.H.; Jang, J.; Hahn, J.-S. Activation of Haa1 and War1 transcription factors by differential binding of weak acid anions in Saccharomyces cerevisiae. Nucleic Acids Res. 2019, 47, 1211–1224, doi:10.1093/nar/gky1188.

16. Simpson-Lavy, K.; Kupiec, M. Carbon Catabolite Repression in Yeast is Not Limited to Glucose. Sci. Rep. 2019, 9, 6491, doi:10.1038/s41598-019-43032-w.

17. Smits, H.; Smits, G.J.; Postma, P.W.; Walsh, M.C.; van Dam, K. High-affinity glucose uptake in Saccharomyces cerevisiae is not dependent on the presence of glucose-phosphorylating enzymes. Yeast 1996, 12, 439–447, doi:10.1002/(SICI)1097-0061(199604)12:5<439:AID-YEA925>3.0.CO;2-W.

18. Nogae, I.; Johnston, M. Isolation and characterization of the ZWF1 gene of Saccharomyces cerevisiae, encoding glucose-6-phosphate dehydrogenase. Gene 1990, 96, 161–169, doi:10.1016/0378-1119(90)90248-p.

19. Tökés-Füzesi, M.; Bedwell, D.M.; Repa, I.; Sipos, K.; Sümegi, B.; Rab, A.; Miseta, A. Hexose phosphorylation and the putative calcium channel component Mid1p are required for the hexose-induced transient elevation of cytosolic calcium response in Saccharomyces cerevisiae. Molecular Microbiology 2002, 44, 1299–1308, doi:10.1046/j.1365-2958.2002.02956.x.

20. Hong, M.-P.; Vu, K.; Bautos, J.; Gelli, A. Cch1 restores intracellular Ca2+ in fungal cells during endoplasmic reticulum stress. Journal of Biological Chemistry 2010, 285, 10951–10958, doi:10.1074/jbc.M109.056218.

21. Locke, E.G.; Bonilla, M.; Liang, L.; Takita, Y.; Cunningham, K.W. A homolog of voltage-gated Ca(2+) channels stimulated by depletion of secretory Ca(2+) in yeast. Mol. Cell. Biol. 2000, 20, 6686–6694, doi:10.1128/MCB.20.18.6686-6694.2000.

22. Batiza, A.F.; Schulz, T.; Masson, P.H. Yeast respond to hypotonic shock with a calcium pulse. J. Biol. Chem. 1996, 271, 23357–23362, doi:10.1074/jbc.271.38.23357.

23. Paidhungat, M.; Garrett, S. A homolog of mammalian, voltage-gated calcium channels mediates yeast pheromone-stimulated Ca2+ uptake and exacerbates the cdc1(Ts) growth defect. Mol. Cell. Biol. 1997, 17, 6339–6347, doi:10.1128/MCB.17.11.6339.

24. Miseta, A.; Kellermayer, R.; Aiello, D.P.; Fu, L.; Bedwell, D.M. The vacuolar Ca2+/H+ exchanger Vcx1p/Hum1p tightly controls cytosolic Ca2+ levels in S. cerevisiae. FEBS Lett. 1999, 451, 132– 136, doi:10.1016/s0014-5793(99)00519-0.

25. Miseta, A.; Fu, L.; Kellermayer, R.; Buckley, J.; Bedwell, D.M. The Golgi apparatus plays a significant role in the maintenance of Ca2+ homeostasis in the vps33Delta vacuolar biogenesis mutant of Saccharomyces cerevisiae. J. Biol. Chem. 1999, 274, 5939–5947, doi:10.1074/jbc.274.9.5939.

26. Halachmi, D.; Eilam, Y. Calcium homeostasis in yeast cells exposed to high concentrations of calcium. Roles of vacuolar H(+)-ATPase and cellular ATP. FEBS Lett. 1993, 316, 73–78, doi:10.1016/0014-5793(93)81739-m.

27. Groppi, S.; Belotti, F.; Brandão, R.L.; Martegani, E.; Tisi, R. Glucose-induced calcium influx in budding yeast involves a novel calcium transport system and can activate calcineurin. Cell Calcium 2011, 49, 376–386, doi:10.1016/j.ceca.2011.03.006.

28. Peiter, E.; Fischer, M.; Sidaway, K.; Roberts, S.K.; Sanders, D. The Saccharomyces cerevisiae Ca2+ channel Cch1pMid1p is essential for tolerance to cold stress and iron toxicity. 0014-5793 2005, 579, 5697–5703, doi:10.1016/j.febslet.2005.09.058.

29. Viladevall, L.; Serrano, R.; Ruiz, A.; Domenech, G.; Giraldo, J.; Barceló, A.; Ariño, J. Characterization of the calcium-mediated response to alkaline stress in Saccharomyces cerevisiae. J. Biol. Chem. 2004, 279, 43614–43624, doi:10.1074/jbc.M403606200.

30. Denis, V.; Cyert, M.S. Internal Ca(2+) release in yeast is triggered by hypertonic shock and mediated by a TRP channel homologue. J Cell Biol 2002, 156, 29–34, doi:10.1083/jcb.200111004.

31. Li, X.; Qian, J.; Wang, C.; Zheng, K.; Ye, L.; Fu, Y.; Han, N.; Bian, H.; Pan, J.; Wang, J.;, et al. Regulating cytoplasmic calcium homeostasis can reduce aluminum toxicity in yeast. PLOS ONE 2011, 6, e21148, doi:10.1371/journal.pone.0021148.

32. Ruta, L.L.; Popa, V.C.; Nicolau, I.; Danet, A.F.; Iordache, V.; Neagoe, A.D.; Farcasanu, I.C. Calcium signaling mediates the response to cadmium toxicity in Saccharomyces cerevisiae cells. FEBS Lett. 2014, 588, 3202–3212, doi:10.1016/j.febslet.2014.07.001.

33. Muller, E.M.; Locke, E.G.; Cunningham, K.W. Differential regulation of two Ca(2+) influx systems by pheromone signaling in Saccharomyces cerevisiae. Genetics 2001, 159, 1527–1538, doi:10.1093/genetics/159.4.1527.

34. The intracellular dissipation of cytosolic calcium following glucose re-addition to carbohydrate depleted Saccharomyces cerevisiae, 2004.

35. Nakajima-Shimada, J.; Iida, H.; Tsuji, F.I.; Anraku, Y. Monitoring of intracellular calcium in Saccharomyces cerevisiae with an apoaequorin cDNA expression system. Proc. Natl. Acad. Sci. U. S. A. 1991, 88, 6878–6882, doi:10.1073/pnas.88.15.6878.

36. Pittman, J.K. Vacuolar Ca(2+) uptake. Cell Calcium 2011, 50, 139–146, doi:10.1016/j.ceca.2011.01.004.

37. D’hooge, P.; Coun, C.; van Eyck, V.; Faes, L.; Ghillebert, R.; Mariën, L.; Winderickx, J.; Callewaert, G. Ca(2+) homeostasis in the budding yeast Saccharomyces cerevisiae: Impact of ER/Golgi Ca(2+) storage. Cell Calcium 2015, 58, 226–235, doi:10.1016/j.ceca.2015.05.004.

38. Thomas, D.; Cherest, H.; Surdin-Kerjan, Y. Identification of the structural gene for glucose-6-phosphate dehydrogenase in yeast. Inactivation leads to a nutritional requirement for organic sulfur. EMBO J. 1991, 10, 547–553, doi:10.1002/j.1460-2075.1991.tb07981.x.

39. Aiello, D.P.; Fu, L.; Miseta, A.; Bedwell, D.M. Intracellular glucose 1-phosphate and glucose 6-phosphate levels modulate Ca2+ homeostasis in Saccharomyces cerevisiae. J. Biol. Chem. 2002, 277, 45751–45758, doi:10.1074/jbc.M208748200.

40. Csutora, P.; Strassz, A.; Boldizsár, F.; Németh, P.; Sipos, K.; Aiello, D.P.; Bedwell, D.M.; Miseta, A. Inhibition of phosphoglucomutase activity by lithium alters cellular calcium homeostasis and signaling in Saccharomyces cerevisiae. American Journal of Physiology-Cell Physiology 2005, 289, C58–C67, doi:10.1152/ajpcell.00464.2004.

41. Espeso, E.A. The CRaZy Calcium Cycle. Adv. Exp. Med. Biol. 2016, 892, 169–186, doi:10.1007/978-3-319-25304-6_7.

42. Spasskaya, D.S.; Karpov, D.S.; Mironov, A.S.; Karpov, V.L. Transcription factor Rpn4 promotes a complex antistress response in Saccharomyces cerevisiae cells exposed to methyl methanesulfonate. Mol Biol 2014, 48, 141–149, doi:10.1134/S0026893314010130.

43. Ratnakumar, S.; Kacherovsky, N.; Arms, E.; Young, E.T. Snf1 controls the activity of adr1 through dephosphorylation of Ser230. Genetics 2009, 182, 735–745, doi:10.1534/genetics.109.103432.

44. Wang, Y.; Pierce, M.; Schneper, L.; Güldal, C.G.; Zhang, X.; Tavazoie, S.; Broach, J.R. Ras and Gpa2 mediate one branch of a redundant glucose signaling pathway in yeast. PLoS Biol. 2004, 2, E128, doi:10.1371/journal.pbio.0020128.

45. Cannon, J.F.; Tatchell, K. Characterization of Saccharomyces cerevisiae genes encoding subunits of cyclic AMP-dependent protein kinase. Mol. Cell. Biol. 1987, 7, 2653–2663, doi:10.1128/MCB.7.8.2653.

46. Kataoka, T.; Powers, S.; McGill, C.; Fasano, O.; Strathern, J.; Broach, J.; Wigler, M. Genetic analysis of yeast RAS1 and RAS2 genes. Cell 1984, 37, 437–445, doi:10.1016/0092-8674(84)90374-x.

47. Montllor-Albalate, C.; Kim, H.; Thompson, A.E.; Jonke, A.P.; Torres, M.P.; Reddi, A.R. Sod1 integrates oxygen availability to redox regulate NADPH production and the thiol redoxome. Proc. Natl. Acad. Sci. U. S. A. 2022, 119, doi:10.1073/pnas.2023328119.

48. Fu, L.; Miseta, A.; Hunton, D.; Marchase, R.B.; Bedwell, D.M. Loss of the major isoform of phosphoglucomutase results in altered calcium homeostasis in Saccharomyces cerevisiae. J. Biol. Chem. 2000, 275, 5431–5440, doi:10.1074/jbc.275.8.5431.

49. Heinisch, J. Isolation and characterization of the two structural genes coding for phosphofructokinase in yeast. Mol. Gen. Genet. 1986, 202, 75–82, doi:10.1007/BF00330520.

50. Breitenbach-Schmitt, I.; Schmitt, H.D.; Heinisch, J.; Zimmermann, F.K. Genetic and physiological evidence for the existence of a second glycolytic pathway in yeast parallel to the phosphofructokinase-aldolase reaction sequence. Mol. Gen. Genet. 1984, 195, 536–540, doi:10.1007/BF00341459.

51. Klinder, A.; Kirchberger, J.; Edelmann, A.; Kopperschläger, G. Assembly of phosphofructokinase-1 fromSaccharomyces cerevisiae in extracts of single-deletion mutants. Yeast 1998, 14, 323–334, doi:10.1002/(SICI)1097-0061(19980315)14:4<323:AID-YEA223>3.0.CO;2-W.

52. Chan, C.-Y.; Parra, K.J. Yeast phosphofructokinase-1 subunit Pfk2p is necessary for pH homeostasis and glucose-dependent vacuolar ATPase reassembly. J. Biol. Chem. 2014, 289, 19448–19457, doi:10.1074/jbc.M114.569855.

53. Heux, S.; Cadiere, A.; Dequin, S. Glucose utilization of strains lacking PGI1 and expressing a transhydrogenase suggests differences in the pentose phosphate capacity among Saccharomyces cerevisiae strains. FEMS Yeast Res. 2008, 8, 217–224, doi:10.1111/j.1567-1364.2007.00330.x.

54. Boles, E.; Liebetrau, W.; Hofmann, M.; Zimmermann, F.K. A family of hexosephosphate mutases in Saccharomyces cerevisiae. Eur. J. Biochem. 1994, 220, 83–96, doi:10.1111/j.1432-1033.1994.tb18601.x.

55. Stathopoulos, A.M.; Cyert, M.S. Calcineurin acts through the CRZ1/TCN1-encoded transcription factor to regulate gene expression in yeast. Genes Dev. 1997, 11, 3432–3444, doi:10.1101/gad.11.24.3432.

56. Puigpinós, J.; Casas, C.; Herrero, E. Altered intracellular calcium homeostasis and endoplasmic reticulum redox state in Saccharomyces cerevisiae cells lacking Grx6 glutaredoxin. Mol. Biol. Cell 2015, 26, 104–116, doi:10.1091/mbc.E14-06-1137.

57. Cunningham, K.W.; Fink, G.R. Calcineurin-dependent growth control in Saccharomyces cerevisiae mutants lacking PMC1, a homolog of plasma membrane Ca2+ ATPases. J Cell Biol 1994, 124, 351–363, doi:10.1083/jcb.124.3.351.

58. Colinet, A.-S.; Sengottaiyan, P.; Deschamps, A.; Colsoul, M.-L.; Thines, L.; Demaegd, D.; Duchêne, M.-C.; Foulquier, F.; Hols, P.; Morsomme, P. Yeast Gdt1 is a Golgi-localized calcium transporter required for stress-induced calcium signaling and protein glycosylation. Sci. Rep. 2016, 6, 24282, doi:10.1038/srep24282.

59. Vashist, S.; Frank, C.G.; Jakob, C.A.; Ng, D.T.W. Two distinctly localized p-type ATPases collaborate to maintain organelle homeostasis required for glycoprotein processing and quality control. Mol. Biol. Cell 2002, 13, 3955–3966, doi:10.1091/mbc.02-06-0090.

60. Zhao, Y.; Du, J.; Zhao, G.; Jiang, L. Activation of calcineurin is mainly responsible for the calcium sensitivity of gene deletion mutations in the genome of budding yeast. Genomics 2013, 101, 49–56, doi:10.1016/j.ygeno.2012.09.005.

61. Ma, T.-Y.; Deprez, M.-A.; Callewaert, G.; Winderickx, J. Coordinated glucose-induced Ca2+ and pH responses in yeast Saccharomyces cerevisiae. Cell Calcium 2021, 100, 102479, doi:10.1016/j.ceca.2021.102479.

62. Simpson-Lavy, K.J.; Kupiec, M. Regulation of yeast Snf1 (AMPK) by a polyhistidine containing pH sensing module. iScience 2022, 25, 105083, doi:10.1016/j.isci.2022.105083.

63. Geva, Y.; Crissman, J.; Arakel, E.C.; Gómez-Navarro, N.; Chuartzman, S.G.; Stahmer, K.R.; Schwappach, B.; Miller, E.A.; Schuldiner, M. Two novel effectors of trafficking and maturation of the yeast plasma membrane H+ -ATPase. Traffic 2017, 18, 672–682, doi:10.1111/tra.12503.

64. Cunningham, K.W.; Fink, G.R. Calcineurin inhibits VCX1-dependent H+/Ca2+ exchange and induces Ca2+ ATPases in Saccharomyces cerevisiae. Mol. Cell. Biol. 1996, 16, 2226–2237, doi:10.1128/MCB.16.5.2226.

65. Masuda, C.A.; Xavier, M.A.; Mattos, K.A.; Galina, A.; Montero-Lomeli, M. Phosphoglucomutase is an in vivo lithium target in yeast. J. Biol. Chem. 2001, 276, 37794–37801, doi:10.1074/jbc.M101451200.

66. Newcomb, L.L.; Diderich, J.A.; Slattery, M.G.; Heideman, W. Glucose regulation of Saccharomyces cerevisiae cell cycle genes. Eukaryot. Cell 2003, 2, 143–149, doi:10.1128/EC.2.1.143-149.2003.

67. Ruiz, A.; Serrano, R.; Ariño, J. Direct regulation of genes involved in glucose utilization by the calcium/calcineurin pathway. J. Biol. Chem. 2008, 283, 13923–13933, doi:10.1074/jbc.M708683200.

68. Simpson-Lavy, K.; Xu, T.; Johnston, M.; Kupiec, M. The Std1 Activator of the Snf1/AMPK Kinase Controls Glucose Response in Yeast by a Regulated Protein Aggregation. Mol. Cell 2017, 68, 1120–1133.e3, doi:10.1016/j.molcel.2017.11.016.

69. Qu, Y.; Jiang, J.; Liu, X.; Wei, P.; Yang, X.; Tang, C. Cell Cycle Inhibitor Whi5 Records Environmental Information to Coordinate Growth and Division in Yeast. Cell Rep. 2019, 29, 987–994.e5, doi:10.1016/j.celrep.2019.09.030.

70. Stockwell, S.R.; Landry, C.R.; Rifkin, S.A. The yeast galactose network as a quantitative model for cellular memory. Mol. Biosyst. 2015, 11, 28–37, doi:10.1039/c4mb00448e.

71. Velivela, S.D.; Kane, P.M. Compensatory Internalization of Pma1 in V-ATPase Mutants in Saccharomyces cerevisiae Requires Calcium- and Glucose-Sensitive Phosphatases. Genetics 2018, 208, 655–672, doi:10.1534/genetics.117.300594.

72. Kim, Y.J.; Lee, M.K.; Kim, U.; Lee, J.-M.; Hsieh, Y.S.; Seol, G.H. Lavandula angustifolia Mill. inhibits high glucose and nicotine-induced Ca2+ influx in microglia and neuron-like cells via two distinct mechanisms. Biomed. Pharmacother. 2024, 177, 117062, doi:10.1016/j.biopha.2024.117062.

73. Delgadillo-Silva, L.F.; Tasöz, E.; Singh, S.P.; Chawla, P.; Georgiadou, E.; Gompf, A.; Rutter, G.A.; Ninov, N. Optogenetic β cell interrogation in vivo reveals a functional hierarchy directing the Ca2+ response to glucose supported by vitamin B6. Sci. Adv. 2024, 10, eado4513, doi:10.1126/sciadv.ado4513.

74. Babenko, V.A.; Varlamova, E.G.; Saidova, A.A.; Turovsky, E.A.; Plotnikov, E.Y. Lactate protects neurons and astrocytes against ischemic injury by modulating Ca2+ homeostasis and inflammatory response. FEBS J. 2024, 291, 1684–1698, doi:10.1111/febs.17051.

75. Zaborska, K.E.; Dadi, P.K.; Dickerson, M.T.; Nakhe, A.Y.; Thorson, A.S.; Schaub, C.M.; Graff, S.M.; Stanley, J.E.; Kondapavuluru, R.S.; Denton, J.S.;, et al. Lactate activation of α-cell KATP channels inhibits glucagon secretion by hyperpolarizing the membrane potential and reducing Ca2+ entry. Mol. Metab. 2020, 42, 101056, doi:10.1016/j.molmet.2020.101056.

76. Raghav, D.; Shukla, S.; Jadiya, P. Mitochondrial calcium signaling in non-neuronal cells: Implications for Alzheimer’s disease pathogenesis. Biochim. Biophys. Acta Mol. Basis Dis. 2024, 1870, 167169, doi:10.1016/j.bbadis.2024.167169.

77. Mitaishvili, E.; Feinsod, H.; David, Z.; Shpigel, J.; Fernandez, C.; Sauane, M.; La Parra, C. de. The Molecular Mechanisms behind Advanced Breast Cancer Metabolism: Warburg Effect, OXPHOS, and Calcium. Front. Biosci. (Landmark Ed) 2024, 29, 99, doi:10.31083/j.fbl2903099.

78. Saint-Prix, F.; Bönquist, L.; Dequin, S. Functional analysis of the ALD gene family of Saccharomyces cerevisiae during anaerobic growth on glucose: the NADP+-dependent Ald6p and Ald5p isoforms play a major role in acetate formation. Microbiology (Reading) 2004, 150, 2209–2220, doi:10.1099/mic.0.26999-0.

79. Baker Brachmann, C.; Davies, A.; Cost, G.J.; Caputo, E.; Li, J.; Hieter, P.; Boeke, J.D. Designer deletion strains derived fromSaccharomyces cerevisiae S288C: A useful set of strains and plasmids for PCR-mediated gene disruption and other applications. Yeast 1998, 14, 115–132, doi:10.1002/(SICI)1097-0061(19980130)14:2<115:AID-YEA204>3.0.CO;2-2.

80. Ma, H.; Kunes, S.; Schatz, P.J.; Botstein, D. Plasmid construction by homologous recombination in yeast. Gene 1987, 58, 201–216, doi:10.1016/0378-1119(87)90376-3.

81. Nayak, V.; Zhao, K.; Wyce, A.; Schwartz, M.F.; Lo, W.-S.; Berger, S.L.; Marmorstein, R. Structure and dimerization of the kinase domain from yeast Snf1, a member of the Snf1/AMPK protein family. Structure 2006, 14, 477–485, doi:10.1016/j.str.2005.12.008.

82. Orlova, M.; Barrett, L.; Kuchin, S. Detection of endogenous Snf1 and its activation state: application to Saccharomyces and Candida species. Yeast 2008, 25, 745–754, doi:10.1002/yea.1628.

