## Supplementary figures and images for "Calcium signaling is a universal carbon source signal transducer and effects an ionic memory of past carbon sources"

### Figure S1.tif

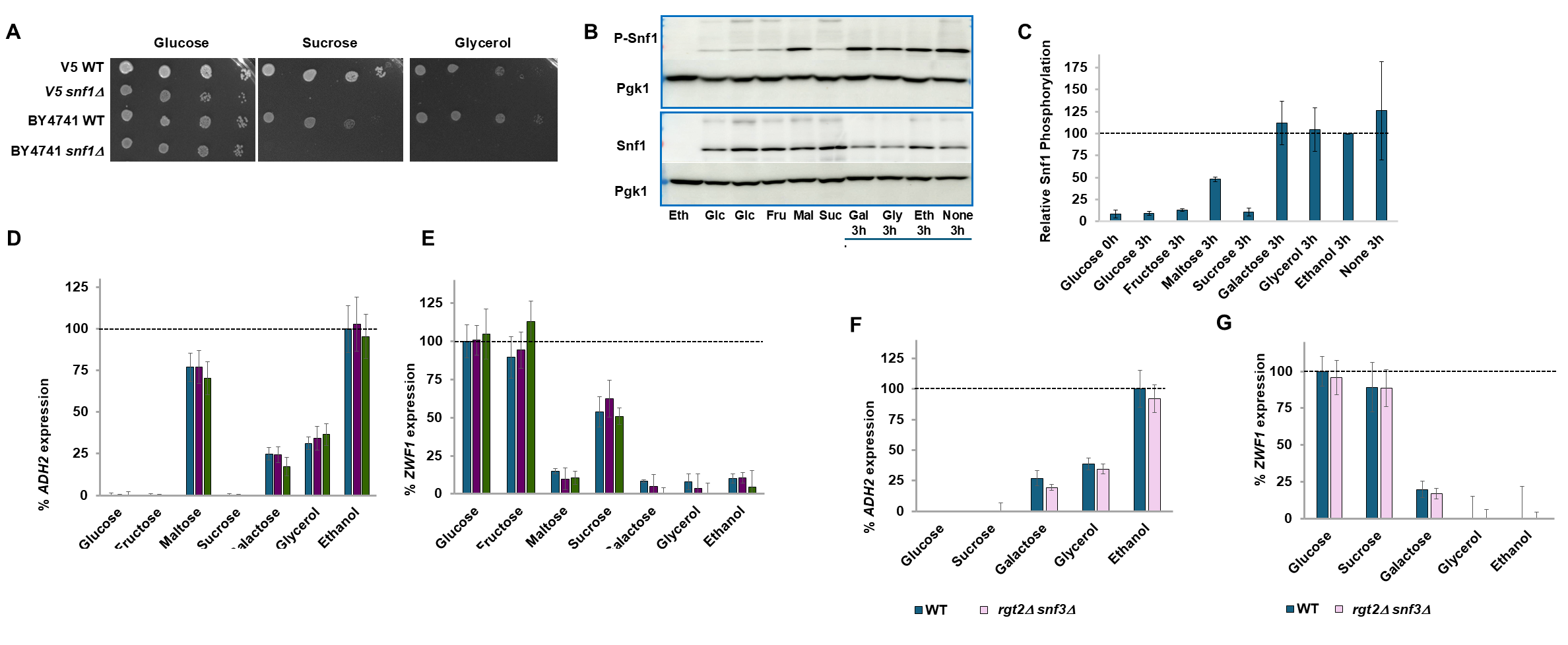

### Figure S2.tif

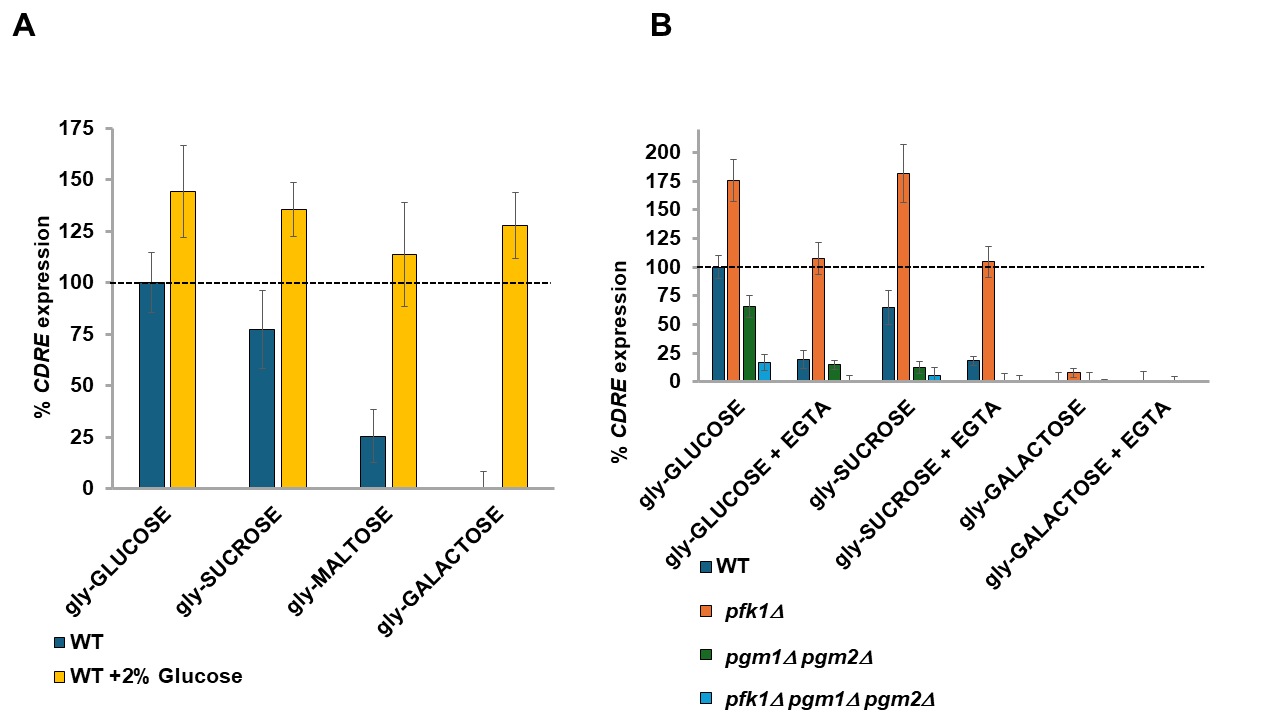

### Figure S3.tif

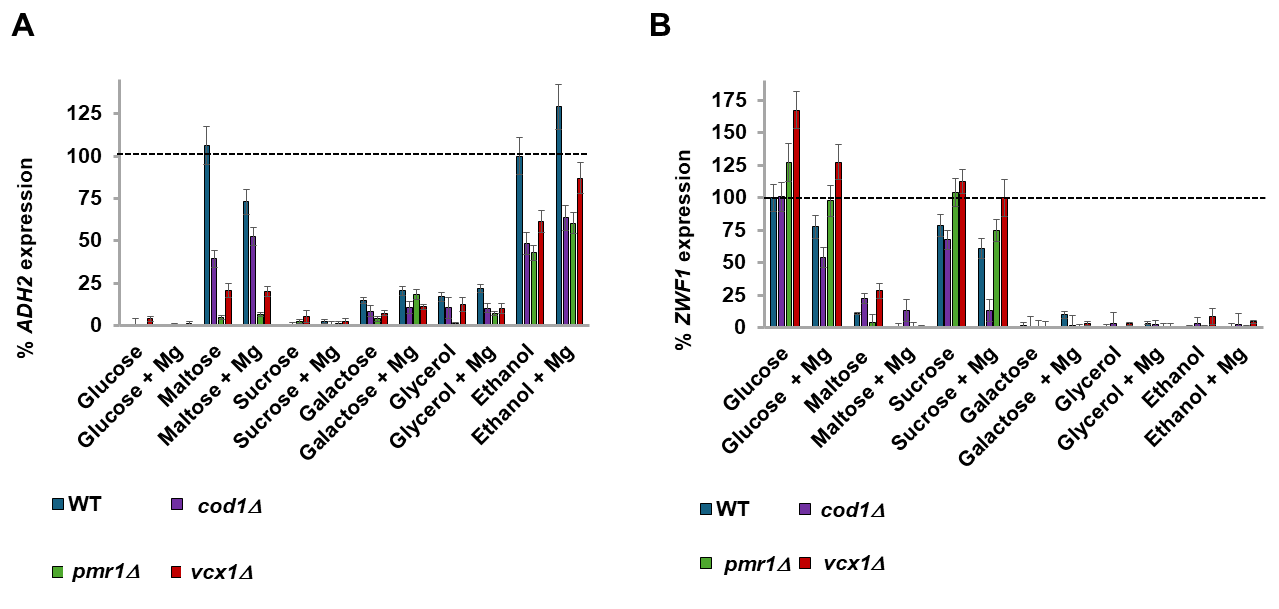

### Figure S4.tif

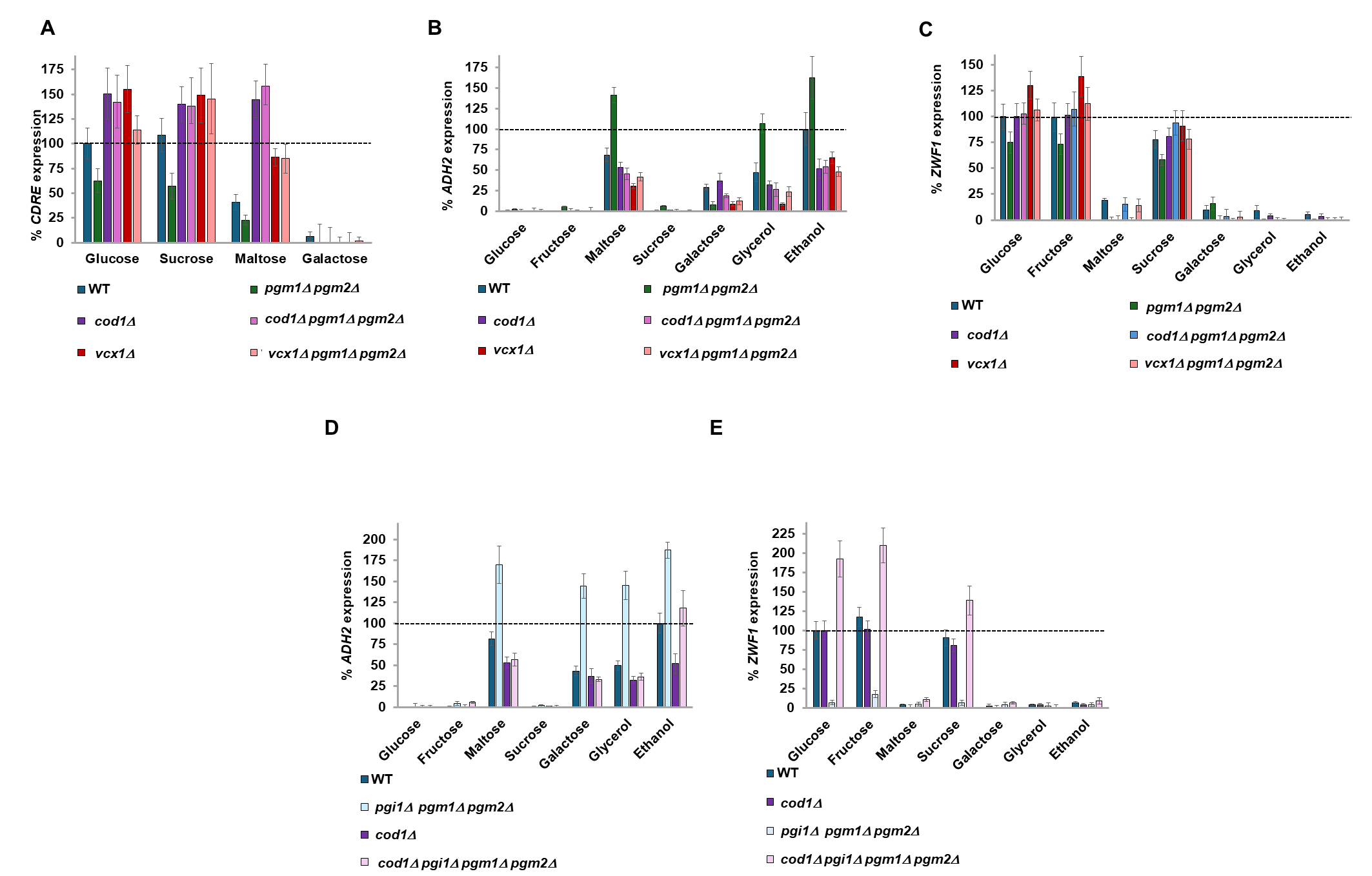

### Figure S5.tif

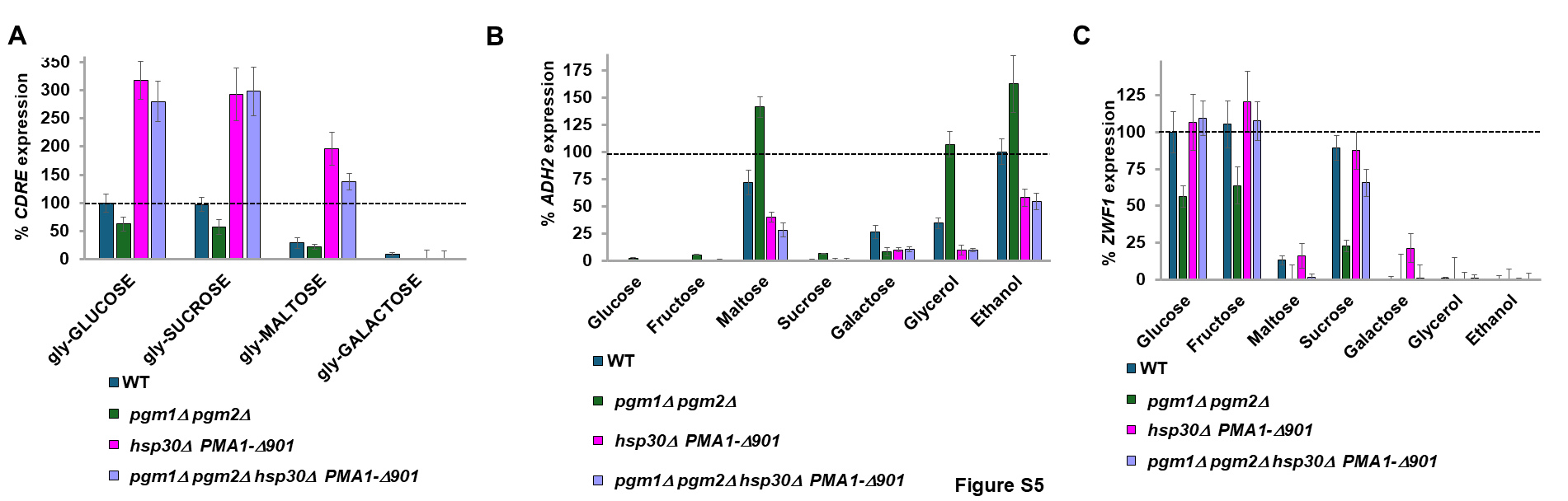

### Figure S6.tif

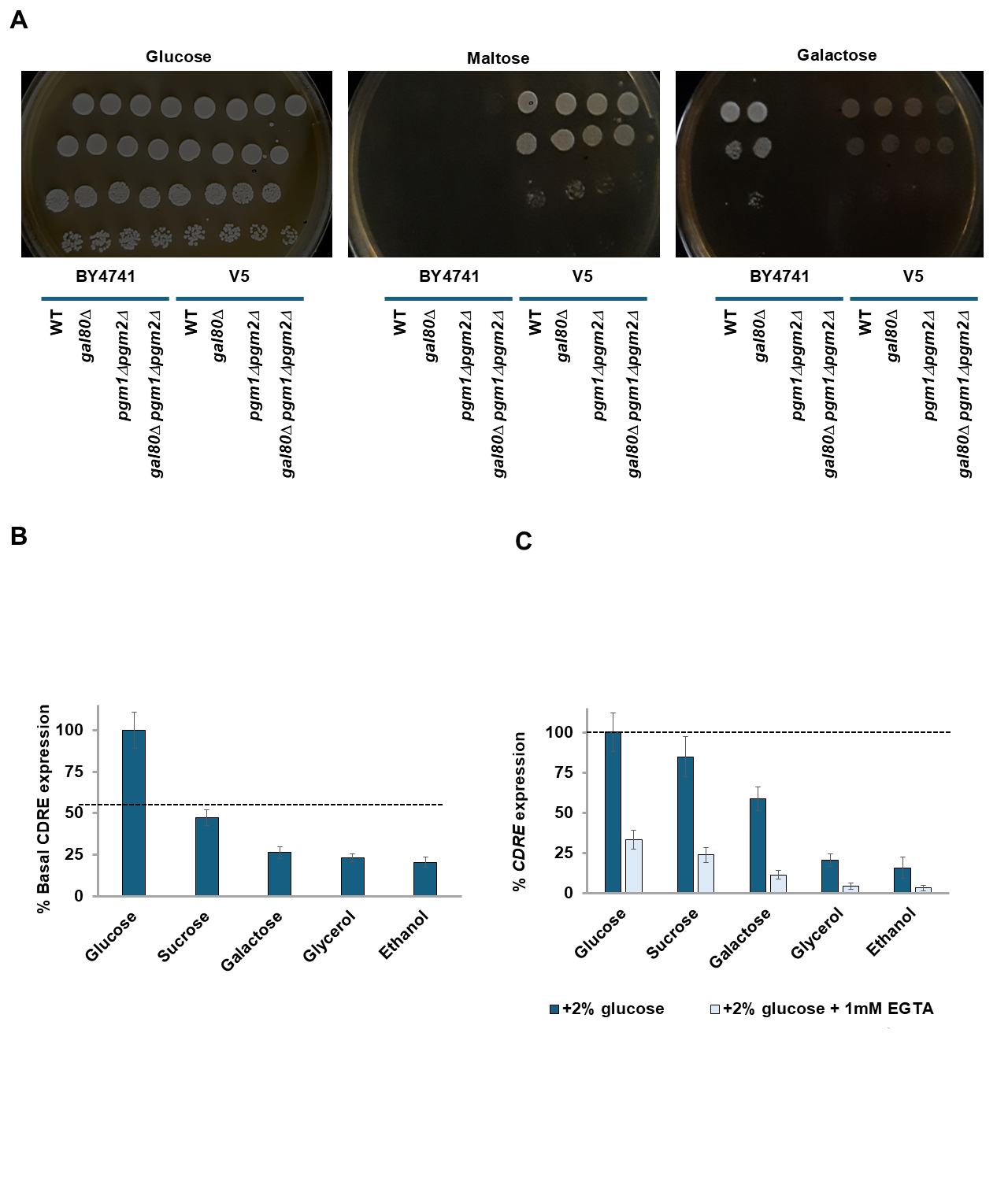
